# The effect of Wnt signaling on the localization, molecular size and activity of the beta-catenin destruction complex in vivo

**DOI:** 10.1101/177790

**Authors:** Kristina N. Schaefer, Shiping Zhang, Teresa T. Bonello, Clara E. Williams, Daniel J. McKay, Mark Peifer

## Abstract

Wnt signaling provides a paradigm for cell-cell signals that regulate embryonic development and stem cell homeostasis and are inappropriately activated in cancers. Our current outline of Wnt signaling focuses around several key players. The tumor suppressors APC and Axin form the core of the multiprotein destruction complex, which targets the Wnt-effector beta-catenin for phosphorylation, ubiquitination and destruction. However, mechanisms underlying destruction complex function and those by which Wnt signaling inactivates it remain much less clear. Based on work in cultured cells, we hypothesize the destruction complex is a supermolecular entity that self-assembles by Axin and APC polymerization, and that regulating complex assembly and dynamics underlie function. We took these insights into the *Drosophila* embryonic epidermis, a premier model of Wnt signaling. Combining biochemistry, genetic tools to manipulate Axin and APC2 levels, advanced imaging and molecule counting, we defined destruction complex assembly, stoichiometry, and localization in vivo, and its downregulation in response to Wnt signaling. Our findings challenge and revise current models of destruction complex function. Endogenous Axin and APC2 proteins accumulate at roughly similar levels, countering the accepted dogma that Axin accumulates at much lower levels. By expressing Axin:GFP at near endogenous levels we found Axin assembles into large cytoplasmic complexes containing tens to hundreds of Axin proteins. Wnt signals trigger complex recruitment to the membrane, while diffuse cytoplasmic Axin levels increase, suggesting slowed assembly. Manipulating Axin or APC2 levels had no effect on destruction complex activity when Wnt signals were absent, but, surprisingly, had opposite effects on the destruction complex when Wnt signals were present. Elevating Axin made the complex resistant to inactivation, while elevating APC2 levels enhanced inactivation. Our data suggest both absolute levels and the ratio of these two core components affect destruction complex function, supporting models in which competition among Axin partners determines destruction complex activity.

**Author Summary:** Cell-cell communication is critical for cells to choose fates during embryonic development and often goes wrong in diseases like cancer. The Wnt cell signaling pathway provides a superb example. Mutations in negative regulators like the proteins APC and Axin take the brakes off cell proliferation and thus contribute to colon cancer. We study how APC, Axin and their protein partners keep cell signaling off, and how cell-to-cell Wnt signals reverse this. We use the fruit fly embryo, combining biochemical, and genetic tools with advanced microscopy. We found that APC2 and Axin proteins are present in cells in similar numbers, challenging the previous dogma. We further find that the ability of Wnt signaling to turn off this negative regulatory machine is influenced both by the levels of Axin and APC2 and by the ratio of their protein levels. We also visualize the active destruction complex in the animal, and count the number of Axin proteins in this complex. Finally, we find that Wnt signals have two effects on the destruction complex—recruiting it to the cell’s plasma membrane and reducing its ability to assemble. Based on this, we propose a new model for how this important signaling pathway is regulated.

**Abbreviations:**

ßcat: beta-catenin
APC: adenomatous polyposis coli
GSK3: glycogen synthase kinase-3
CK1: casein kinase 1
Dsh: Dishevelled
DIX domain: Dishevelled/Axin domain
LRP5/6: low-density lipoprotein receptor-related proteins 5/6
GFP: green fluorescent protein
RFP: red fluorescent protein
ROI: rectangular region of interest
TCF: T-cell factor
Wg: Wingless

## Introduction

Cell-cell signaling is critical for cell fate decisions during embryonic development and cell fate maintenance during adult homeostasis. Altered signaling by these same pathways underlies most solid tumors. The Wnt signaling pathway provides a paradigm—it regulates cell fate choice in tissues throughout the body, maintains stem cell identity in many adult tissues, and is inappropriately activated in colorectal and other cancers [1]. Thus, understanding the mechanisms by which signaling occurs and is regulated are key issues for cell, developmental, and cancer biology.

Work in both animal models and cultured mammalian cells provided a broad outline of Wnt signaling and its regulation [2]. The key effector is the transcriptional co-activator βcatenin (βcat; *Drosophila* Armadillo; Arm). In the absence of signaling, βcat is captured by a multiprotein complex called the destruction complex. The scaffold proteins Adenomatous polyposis coli (APC) and Axin bind βcat and present it to the kinases glycogen synthase kinase-3 (GSK3) and casein kinase 1 (CK1). They phosphorylate βcat, creating a binding site for an E3 ubiquitin ligase, thus targeting βcat for proteasomal destruction. When Wnt ligands bind to receptors, the destruction complex is inactivated, allowing ßcat to accumulate, enter the nucleus and act together with the DNA binding protein TCF to transcriptionally activate Wnt-regulated genes.

While this broad outline is well-supported, the underlying mechanisms by which the destruction complex targets ßcat for destruction, and by which Wnt signaling inactivates it remain much less clear. APC was originally viewed as the scaffold around which the destruction complex assembled, but subsequent work revealed that Axin fulfills this function, leaving APC’s molecular role a mystery. Further, while the destruction complex is typically represented in models as a simple four-protein complex, considerable evidence supports the idea that it is a large supermolecular protein assembly, built by self-polymerization of Axin and APC (e.g., [3-6]).

Recent work provided new mechanistic insights into the underlying molecular mechanisms, helping us begin to transform the static, low-resolution textbook model of Wnt signaling into a more dynamic, high resolution view. Super-resolution microscopy of Axin and APC complexes assembled after overexpression in colorectal cancer cells provided the first look inside the active destruction complex. Axin and APC containing “puncta” were resolved into intertwined strands of each protein, presumably assembled by polymerization. Combining this with assessment of APC and Axin dynamics as well as genetic and biochemical dissection of the two proteins provided novel mechanistic insights and a new model. Together with previous work, these data suggest that APC first promotes Axin multimerization, and then, after Axin-mediated βcat phosphorylation, APC undergoes a regulated conformational change that transfers βcat out of the destruction complex to the E3 ligase, and then restarts the catalytic cycle. These data fit with other recent studies suggesting that Wnt signaling does not totally turn off the destruction complex. Instead, the destruction complex remains intact and capable of phosphorylating βcat, but βCat transfer to the E3 ligase is prevented [5,7-9].

However, this work was largely done in cultured cells, which provide a simple place to explore pathway circuitry but do not provide a physiologically relevant situation with all regulatory mechanisms intact. We thus took these insights back into the *Drosophila* embryonic epidermis, arguably the system where our understanding of the roles and regulation of the Wnt pathway is strongest. Stripes of cells in each body segment produce the fly Wnt, Wingless (Wg), creating a field of cells experiencing high, moderate and low levels of Wg signaling. Taking advantage of new genetic approaches and high resolution microscopy, we sought to address several key issues in the field, involving the structure, assembly and stoichiometry of the destruction complex in vivo during normal development, and how it is downregulated by Wnt signaling.

To understand assembly of a complex multiprotein machine, one key issue involves the relative levels of its component parts. Current models of Wnt regulation suggest Axin accumulates at levels <u>dramatically</u> lower than those of other proteins in the destruction complex. This hypothesis is largely based on early work in *Xenopus* oocyte extracts. By adding in known amounts of recombinant Axin and measuring the resulting destruction complex activity, they estimated Axin concentrations were as much as 5000-fold lower than those of APC and other destruction complex proteins. Their mathematical model of Wnt signaling and most subsequent ones are based on these estimates [10,11]. In contrast, recent work in cultured mammalian cells suggests Axin and APC levels are more similar [12]. Thus, defining the relative levels of Axin and APC in tissues undergoing Wnt signaling in vivo is a key issue, and the *Drosophila* embryo provided a superb place to accomplish this end.

Once relative protein levels are defined, different models for the function and regulation of the destruction complex can be tested by varying absolute levels of Axin or APC and their relative ratios to one another. Substantially elevating Axin levels in *Drosophila* embryos strongly inhibits Wnt signaling [13]. Further analyses suggested there is a threshold below which elevating Axin does not alter signaling, since more subtle elevation of Axin levels (2-5 fold) had little effect in *Drosophila* embryos or imaginal discs [14-16] and mutating *tankyrase*, which elevates Axin levels 2-3 fold, does not substantially perturb Wnt signaling [15,17]. In contrast a 9-fold increase in Axin levels inhibited Wnt signaling in imaginal discs [14]. Together these data suggest that there may be a sharp threshold over which increasing Axin levels inhibits Wnt signaling. However, these studies used multiple tissues or systems in parallel, and left the mechanisms underlying the sharp threshold unclear. The *Drosophila* embryo provided a place to assess how altering Axin levels affects cell fate choice, Wnt-target gene expression and ßcat levels in parallel, and to directly compare effects on cells receiving and not receiving Wnt signals. It also offered the opportunity to manipulate APC levels, the other key scaffolding protein in the destruction complex. Whether APC levels are rate-limiting remains a open question, because APC has been viewed as present in substantial excess. The *Drosophila* embryo also allowed us to test effects of varying the Axin:APC ratio, another key parameter of any molecular model.

Finally, to effectively understand destruction complex assembly and function, we need to visualize it directly. Our recent super-resolution imaging of Axin:APC puncta in cultured cells provided the first insights into the internal structure and dynamics of this multiprotein machine, but these experiments involved significant over-expression. The *Drosophila* embryo provided a place to assess whether similar complexes assemble at near endogenous levels. Recent advances in molecular counting technology also offered the possibility of directly assessing the number of Axin proteins assembled in a complex.

Visualizing the destruction complex in the embryo would also allow us to address how Wnt signaling inactivates it. Work in cultured cells led to a model in which Wnt binding the Frizzled:LRP5/6 receptor complex triggers LRP5/6 phosphorylation, and Axin and Dishevelled (Dsh) membrane recruitment [18]. What happens next is disputed, with many events suggested to play a part. For example, some data suggest the destruction complex is disassembled because Dsh competes for Axin [19] or Wnt signaling destabilizes Axin [20]. Interestingly, examining effects of Wg signaling on the destruction complex in *Drosophila* embryos led to starkly divergent conclusions. One group reported that Wg signaling strongly <u>reduced</u> Axin levels, as assessed both by immunofluorescence and immunoblotting [21]. A second, visualizing GFP-tagged Axin, found little or no effect of Wg on Axin levels—instead their data suggested that Wg signaling causes a Dsh-dependent relocalization of Axin from cytoplasmic puncta to the plasma membrane [13]. Finally, a third group reported that Wg signaling initially <u>stabilizes</u> Axin, as assessed by immunofluorescence, increasing both membrane bound and cytoplasmic pools [15,22]. Thus, the effects of Wg signaling on Axin, a key part of the mechanism underlying βcat stabilization, also remain an open question. Our system allowed us to address this issue.

## Results

### *axin* and *APC1/APC2* are transcribed at similar levels

Most current models of Wnt regulation suggest Axin accumulation levels are dramatically lower than those of other destruction complex proteins, thus making destruction complex activity sensitive to very small increases in its levels. However, the literature contains indications that this may not be universally true (e.g. [12]). To better understand how APC2 and Axin levels affect Wnt signaling we directly compared APC2 and Axin levels in *Drosophila* embryos, where we can quantitatively vary protein levels and assess consequences.

We first compared mRNA levels of *Drosophila axin* with those of the genes encoding the two fly APC family proteins, *APC1* and *APC2,* using RNAseq data from staged embryos. In embryos, APC2 plays the predominant role in Wnt regulation during early to mid-embryogenesis ([23-25]; 2-4 hours or 6-8 hours after egg laying, respectively), while APC1 is expressed at low levels early but then becomes prominent in the central nervous system [26,27]. Consistent with this, *APC2* mRNA levels are ∼19x higher than *APC1* during early embryogenesis, and ∼7x higher during mid-embryogenesis (Fragments Per Kilobase of transcript per Million mapped reads (FPKM) 484 versus 26, and FPKM 201 versus 27, respectively). However, at late embryogenesis, as the central nervous system is assembled, *APC1* mRNA levels are now ∼5x more abundant than APC2 (FPKM 120 vs. 23). Since APC2 and APC1 can act redundantly in regulating Wnt signaling [24,25], we compared *axin* mRNA levels with combined mRNA abundance of *APC1* plus *APC2.* Surprisingly, RNAseq reads for *axin* were roughly comparable to those of *APC1* plus *APC2* at three different stages of embryonic development (Fig. 1A), indicating that there are not dramatic differences between APC family members versus that of Axin at the mRNA level.

**Figure 1:**
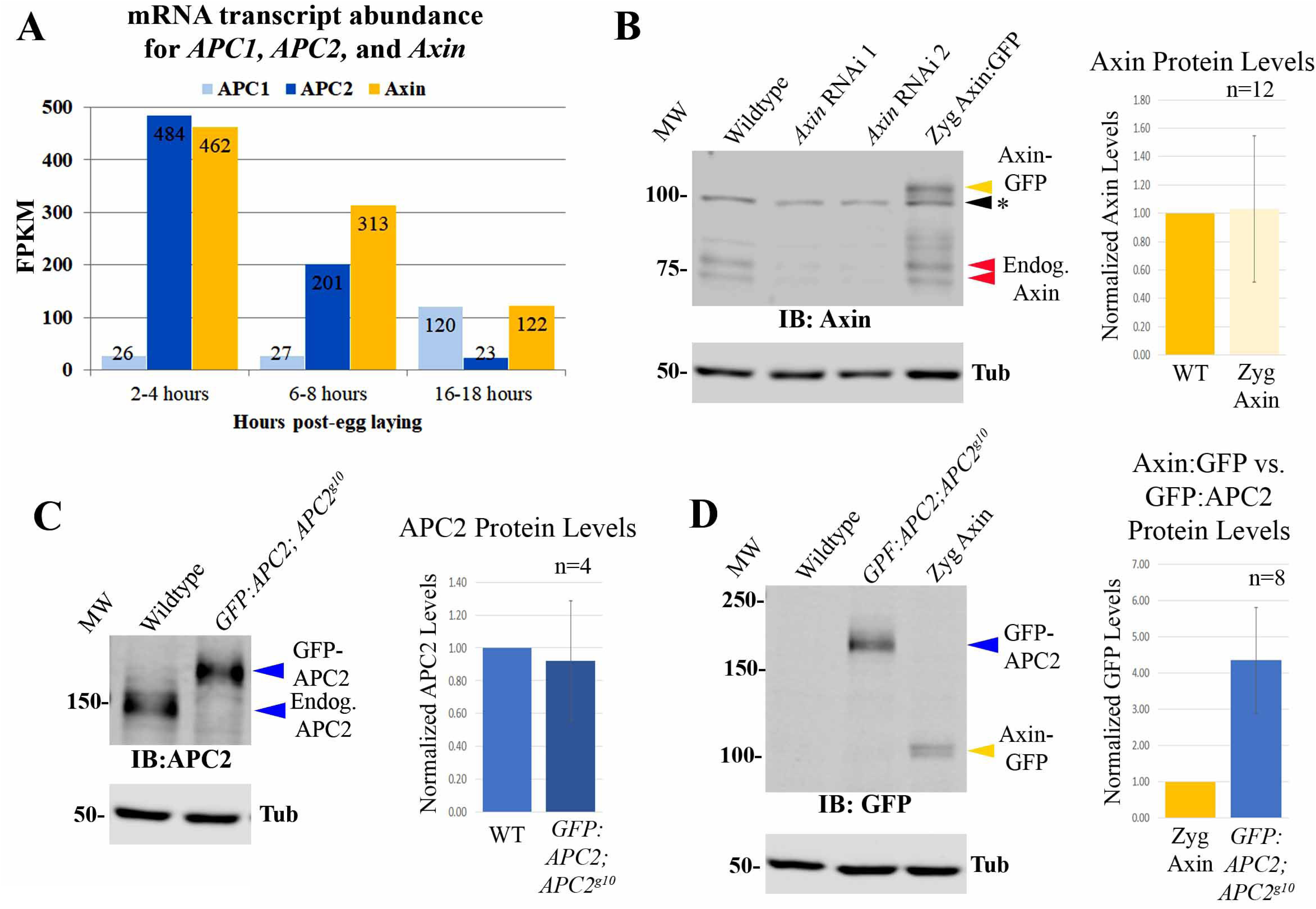
Endogenous APC2 and Axin proteins accumulate at similar levels. (A) mRNA levels (RNAseq) of *APC1* (light blue), *APC2* (blue), and *Axin* (yellow) during *Drosophila* embryogenesis. (B-G) Immunoblots, 4-8hr old *Drosophila* embryos. Tubulin is loading control. n=# of blots quantified (Table S1). (B) Anti-Axin antibody. Endogenous Axin levels versus those in Axin RNAi or Zyg Axin:GFP embryos. Endogenous Axin runs as doublet ∼75kDa (red arrowheads) while GFP Axin runs at ∼105kDa (yellow arrowhead). * = background band. (C) Anti-APC2 antibody. Endogenous APC2 levels versus those of a GFP:APC2 transgene expressed under its endogenous promoter in an *APC2* null (*APC2*^*g10*^) background.(D) Anti-GFP antibody. Relative levels of GFP:APC2 expressed under its endogenous promoter and Zyg Axin:GFP.

### In contrast to current models, Axin and APC2 proteins accumulate at similar levels during early-mid embryogenesis,

These data did not rule out differences in protein translation or stability. To determine if similar transcript levels led to similar protein levels, we compared levels of Axin and APC proteins in early to mid-embryogenesis (4-8 hrs), when APC2 is the predominant family member expressed. Since antibodies to APC2 and Axin may have different affinities, one cannot simply compare antibody-labeled endogenous proteins. To overcome this, we utilized GFP-tagged proteins expressed at near-endogenous levels. This allowed us to compare endogenous versus GFP-tagged Axin, or endogenous versus GFP-tagged APC2 proteins, using antibodies against the endogenous proteins, followed by a comparing GFP-tagged Axin and GFP-tagged APC2 proteins, using anti-GFP antibodies. We used the GAL4-UAS system [28,29] to express Axin:GFP, using the driver that gave the lowest level of Axin:GFP expression (*act5c*-GAL4 provided by male parents). Axin:GFP was expressed at 1.0±0.5 fold that of endogenous Axin, as assessed by immunoblotting with anti-Axin antibodies (Fig. 1B, Table S1). We next used transgenic flies expressing GFP:APC2 under control of the endogenous *APC2* promotor, and crossed it into an *APC2* null mutant background [30]. Using anti-APC2 antibodies, we re-confirmed that *APC2*-driven GFP:APC2 was expressed at essentially the same level as endogenous APC2 (0.9±0.4 fold endogenous APC2; Fig. 1C, Table S1). To complete the comparison, we then compared *APC2*-driven GFP:APC2 to Axin:GFP driven by zygotic *act5c*-GAL4. Immunoblotting with anti-GFP antibodies revealed that GFP:APC2 is expressed ∼4-fold the levels of Axin:GFP (Fig. 1D; 4.3±1.4; Table S1). These three comparisons—endogenous Axin to *act5c*-GAL4 driven Axin:GFP, *act5c*-GAL4 x Axin:GFP to *APC2-*driven GFP:APC2, and *APC2-*driven GFP:APC2 to endogenous APC2—allowed us to make a reasonable estimate of the relative levels of endogenous APC2 to Axin. Our findings reveal that APC2 accumulates at a roughly 5-fold higher level than Axin (4.7±1.4). This is in strong contrast to the 5000-fold difference in accumulation that forms the basis of many current models, but is consistent with the similar levels of mRNAs revealed by RNAseq.

### Developing methods to vary Axin levels during embryogenesis

Another underlying premise of current models of Wnt signaling is that Axin is rate limiting for destruction complex function. Previous experiments in embryos and imaginal discs provide strong support for this, as over-expressing Axin can shut down Wnt signaling. They also suggest this only occurs when Axin levels exceed a specific threshold. Our knowledge of the relative levels of APC2 versus Axin in the *Drosophila* embryonic epidermis allowed us to build on and significantly extend the analysis. We first varied Axin levels systematically, defining the threshold for effects on viability, cell fate and expression of a Wg target gene. We next dug into the underlying mechanism, by examining how different Axin levels affected destruction complex activity and ßcat levels, both in cells receiving and not receiving Wg signals. We then brought APC2 into this picture, examining effects of elevating APC2 levels, and of altering the ratios of Axin to APC2.

To manipulate Axin levels systematically, we used the GAL4-UAS system and several different GAL4 drivers. Four crosses using two different GAL4 drivers provided different levels and timing of Axin over-expression (Fig. S1; Methods; Table S1). *act5c*-GAL4 is expressed both during oogenesis and relatively ubiquitously during embryonic development. The MatGAL4 stock includes two maternally-expressed GAL4 lines, on the second and third chromosomes, which are not expressed zygotically, though maternally expressed GAL4 protein perdures zygotically. 1. By crossing UAS-Axin:GFP females to *act5c*-GAL4/+ males, we achieved lower-level and later elevation of Axin:GFP levels, which was driven by zygotically-expressed GAL4 (hereafter Zyg Axin). 2. By crossing *act5c*-GAL4/+ females to UAS-Axin:GFP males (hereafter Mat/Zyg Axin), we achieved relatively high-level overexpression, which began early due to maternally-contributed GAL4 and continued zygotically. 3. As an alternative for maternal and zygotic over-expression, we created females trans-heterozygous for MatGAL4 and UAS-Axin:GFP (hereafter Mat Axin). 4. To achieve a level of Axin elevation intermediate between that produced by Zyg Axin and Mat Axin, we used MatGAL4 to co-express UAS-Axin:GFP with a second UAS-driven transgene encoding RFP (hereafter Mat RFP&Axin). When two different UAS-driven transgenes are present, this reduces expression of both transgenes. We directly measured protein levels by immunoblotting with antibodies to either GFP or to endogenous Axin.

These four schemes produced an excellent range of Axin expression levels in stage 9 embryos, when Wnt signaling is at its maximum. Zyg Axin effectively tripled normal Axin levels in embryos in which it was expressed (Fig. 2A-C, Table S1; taking into account endogenous Axin and the fact that only 50% of the embryos inherit the GAL4 driver). Mat RFP&Axin resulted in a roughly 4-fold increase, while both Mat/Zyg Axin and Mat Axin led to an 8-9 fold elevation in total Axin levels (Fig. 2A-C, Table S1). When we examined the pattern of Axin:GFP accumulation in embryos, we noted that Mat/Zyg Axin was substantially more variable in expression from cell-cell than MatGAL4 driven Axin:GFP (data not shown). Thus, in most subsequent functional assays we focused on effects of MatGAL4 driven Axin:GFP for high-level overexpression. In addition to differences in expression levels, these lines also differed in timing of Axin:GFP expression (Fig. 2F). Zyg Axin levels started very low (as expected with no maternal GAL4 expression) and continued to rise throughout development. Mat Axin levels started somewhat higher (driven by maternal GAL4), increased during stages 9-11 and then slowly decayed. Mat/Zyg Axin combined features of Zyg Axin and Mat Axin, with initially modest Axin:GFP levels, which continued to rise throughout development. Mat RFP&Axin accumulation followed a similar expression pattern as Mat Axin, but at decreased levels due to the presence of two UAS-driven transgenes (Fig. 2F, right). These tools provided us with the ability to vary Axin levels systematically, and we thus used them to assess how altering Axin levels affects Wg signaling and its regulation, by assessing effects on embryonic viability, cell fate choice, Wg target gene expression, and the levels of the fly homolog of βcat, Arm.

**Figure 2:**
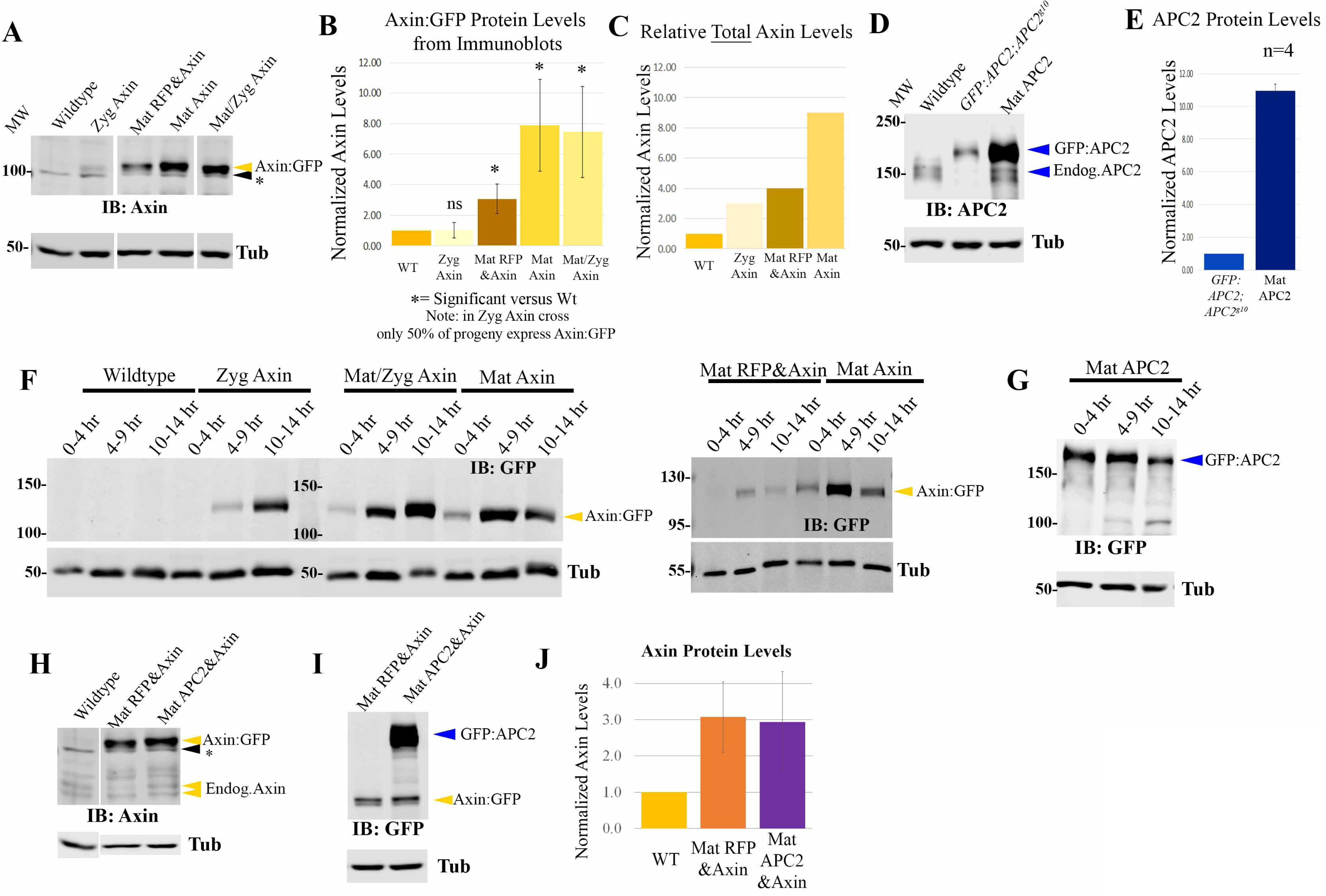
Developing tools to differentially elevate levels of Axin:GFP. (A-E) Immunoblots and quantification, 4-8hr old *Drosophila* embryos. Tubulin is loading control. (A) Anti-Axin antibody. Levels of Axin:GFP when expressed with different GAL4 drivers. *= background band. (B) Quantification, normalized to levels of endogenous Axin. # of blots quantified is in Table S1. (C) Relative levels of total Axin (endogenous Axin + Axin:GFP) accumulation. (D,E) Anti-APC2 antibody. Endogenous APC2, GFP:APC2 expressed via its endogenous promoter, or GFP:APC2 expressed using MatGAL4. (F,G) Immunoblots of Drosophila embryos of the indicated ages, anti-GFP Antibody. Time courses of Axin:GFP (F) or GFP:APC2 (G) accumulation when expressed with different GAL4 drivers. (H) Immunoblot of 4-8hr old *Drosophila* embryos, with anti-Axin antibody. Compares endogenous Axin and Axin:GFP levels in lines expressing both Axin:GFP and a second transgene (RFP or GFP:APC2). From same gel with intervening lanes removed. (I) Same samples stained with an anti-GFP antibody, thus comparing levels of Axin:GFP and GFP:APC2. (J) Quantification of Axin:GFP levels. N=9 blots.

### Over a threshold level Axin inhibits Wg-regulated cell fate choice during embryogenesis

The relatively subtle (3-fold) Axin elevation produced by Zyg Axin did not result in embryonic lethality (Fig. 3A, Table S2; 5% lethality vs. 3% lethality of wildtype controls (UAS-Axin without a GAL4 driver)). We then examined larval cuticles to look for more subtle effects on Wg signaling. Reducing Wg signaling affects cell fate, causing loss of naked cuticle fates and merger of denticle belts—Fig. 3C illustrates the graded series of defects with successively reduced Wg signaling. Most Zyg Axin embryonic cuticles (3-fold increase) were near wildtype (Fig 3B,C, Table S3), though the subtle defects seen suggest subtle reduction of Wg signaling in some embryos. Consistent with this possibility, no hatching Zyg Axin larvae survived to adulthood. The slightly higher level expression of Axin:GFP in Mat RFP&Axin embryos (4-fold increase) and earlier onset of expression led to mild embryonic lethality (32% lethal; Fig. 3A; Table S2), and a larger percentage of embryos had moderate inhibition of Wg signaling, as assessed by cell fate choices (Fig. 3B,C, Table S3). In contrast, higher-level, earlier overexpression of Axin (8-9 fold) led to substantial embryonic lethality—90% lethality of Mat/Zyg Axin and 78% lethality of Mat Axin (Fig. 3A; Table S2). In both crosses, there were two genotypes of embryonic progeny; for Mat/Zyg Axin these differed by whether or not they had a zygotic copy of *act5c*-GAL4 and for Mat Axin by whether they had one or two copies of the UAS-Axin:GFP transgene zygotically (Fig. S1). Cuticle analysis revealed that a significant number of Mat/Zyg Axin and Mat Axin progeny had Wg signaling strongly reduced (Fig. 3B,C, Table S3), but there were variations in the strength of this effect that likely reflect the two different zygotic genotypes in each cross. Thus, there is a relatively sharp threshold (∼4-5 fold endogenous) over which elevating Axin levels led to embryonic lethality due to inhibition of Wg-regulated cell fates.

**Figure 3:**
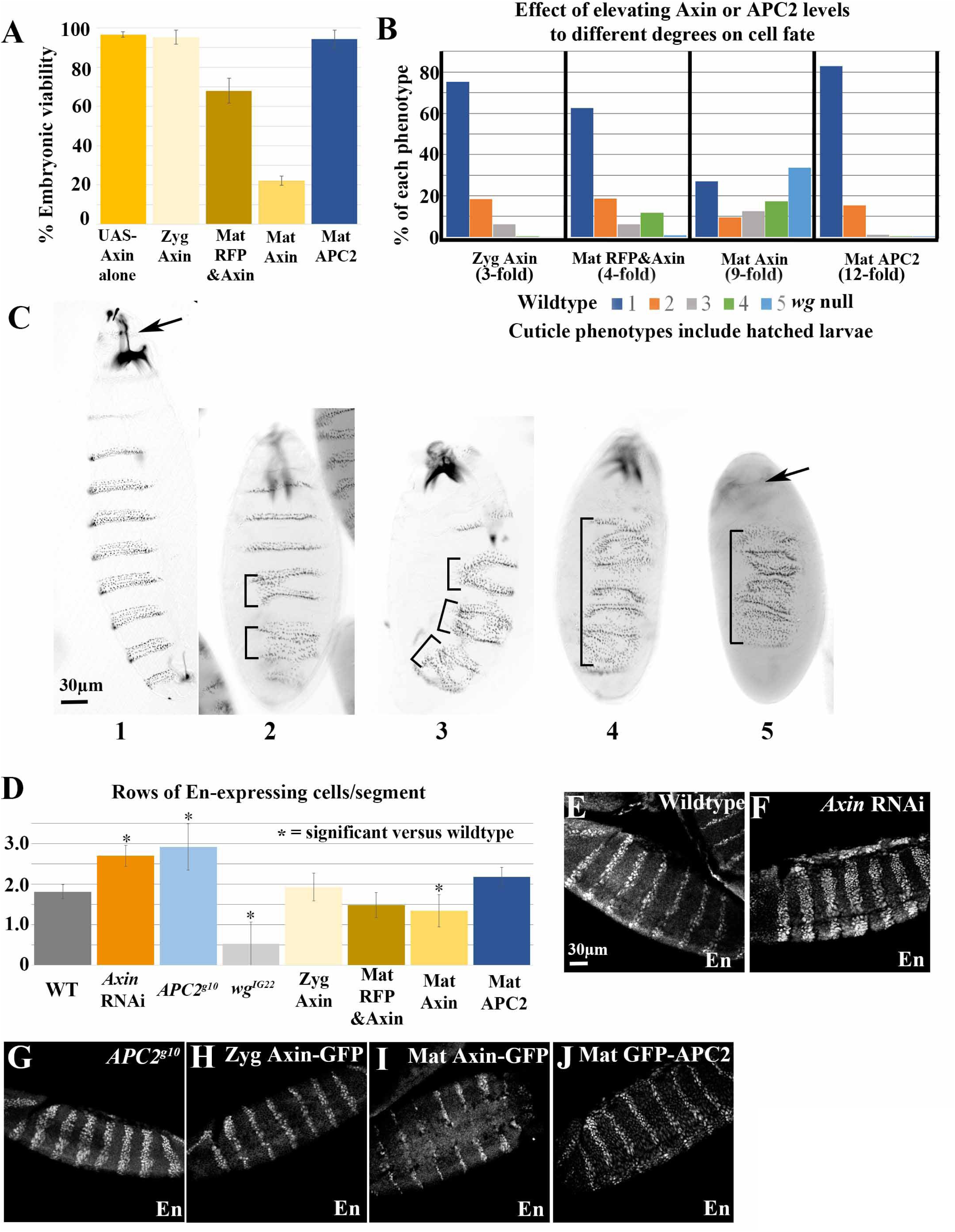
Increasing Axin levels over a threshold inhibits Wg signaling, while increasing APC2 levels does not. (A) Embryonic viability of indicated genotypes. (B,C) Assessing the effect of elevating Axin or APC2 levels on Wg-regulated cell fates. (B) Range of cuticle phenotypes of embryos/larvae of each genotype—since not all genotypes are lethal phenotypes include those of hatched larvae. (C) Representative images of cuticle phenotypes used in B. Anterior to the top. 1: Wildtype. 2: 1-2 merged denticle belts (brackets). 3: 3-4 merged denticle belts. 4: Most denticle belts merged, mouth parts still present. 5: *wg* null phenotype – denticle lawn and no head (arrow). (D) Quantification of number of rows of En-expressing cells per segment. Embryos analyzed: WT-21, AxinRNAi-5, APC2^g10^-5, wg^IG22^ – 14, Zyg Axin-9, Mat RFP&Axin-11, Mat Axin-18, Mat APC2-12. * = p< 0.05 using a one-way ANOVA test. (E-J) Representative images, En expression, as quantified in D. Anterior to the left.

To assess effects of Axin levels on a Wg-regulated target gene, we examined *engrailed* (*en*) expression, using antibodies to its protein product. En usually accumulates in the two most posterior cell rows in each body segment (Fig. 3D,E, Table S4), and maintenance of En expression requires Wg signaling—this is apparent in *wg* mutants in which En stripes are narrowed ([31]; Fig. 3D, Table S4). In contrast, in *APC2*^*g10*^ null mutants or after Axin depletion by RNAi, En expression expands to additional cell rows (Fig. 3D,F,G, Table S4). The 3-fold elevation of Axin levels via ZygGAL4 did not affect En expression (Fig.3D,H, Table S4). In contrast, the 9-fold increase of Axin via MatGAL4 led to partial loss of En expression (Fig. 3D,I, Table S4), though on average this was not as severe as that seen in *wg* mutants. Thus, mildly elevating Axin levels has little effect on Wg regulated cell fates or target genes, but when Axin levels are elevated more than 8-fold, Wg signaling is inhibited.

### Elevating Axin levels over the threshold has no effect on Arm levels in cells <u>not</u> receiving Wg signals, but renders the destruction complex resistant to inactivation by Wg signaling

The primary role of the Axin/APC2 based destruction complex is to regulate levels of the βcat homolog Arm. We thus measured effects of Axin levels on Arm accumulation. Arm has two roles: as part of the cadherin-based cell adhesion complex and as a transcriptional co-activator in the Wnt pathway. Thus, all cells have a pool of Arm at the cortex in adherens junctions. In wildtype, Wg is expressed by one row of cells in each segment, and moves to neighboring cells, resulting in a gradient of Wg signaling across the segment. In cells receiving Wg, the destruction complex is turned down, and Arm accumulates in the cytoplasm and nucleus [32]. The Axin/APC2 based destruction complex also retains Arm in the cytoplasm [30]. Together, these create a gradient of Arm accumulation across the segment, with the highest level of cytoplasmic and nuclear Arm accumulation in the Wg-expressing cells and their immediate neighbors, and gradually decreasing levels of cytoplasmic and nuclear accumulation in cells more distant from the Wg source (Fig. 4A, B diagrammed in B”). In *wg* mutants the destruction complex downregulates Arm in all cells, eliminating the stripes of Arm accumulation [32].

**Figure 4:**
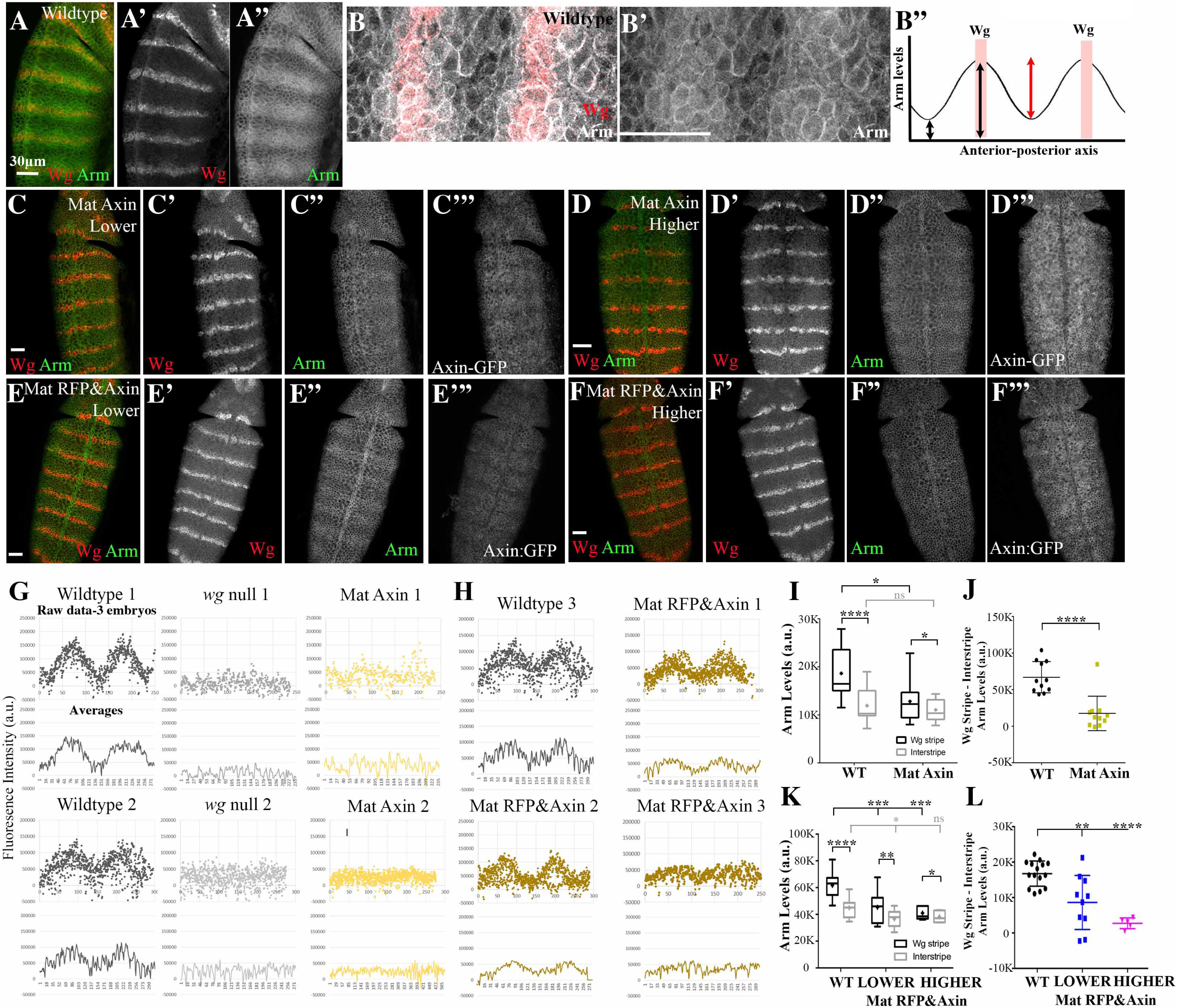
Increasing Axin levels over a threshold reduces the ability of Wg signaling to turn down the destruction complex but has not effect in Wg-Off cells. (A-F) Fixed images, Stage 9 embryos. Anterior to the top. (A,B) Representative images of wildtype embryos. B is a close-up of A. (B”) Diagrammatic illustration of Arm levels across two embryonic segments, illustrating the graded nature of Arm accumulation, and illustrating the parameters we assessed: absolute levels in Wg stripes and in interstripes (black arrows), and the difference between these levels (red arrow)(C,D) Representative images, Mat Axin:GFP embryos with higher or lower levels of Axin:GFP expression, taken from the same slide under the same microscope conditions. Arm accumulation in Wg–stripes is reduced. (E,F) Representative images, Mat RFP&Axin embryos with higher or lower levels of Axin:GFP expression, taken from the same slide under the same microscope conditions. Arm accumulation in Wg –stripes is reduced at higher levels of Axin:GFP expression, but less affected at lower levels. (G,H) Over a threshold, elevating Axin levels flattens the usual graded level of Arm accumulation across each embryonic segment. Plots of Arm accumulation over 2 segments for each genotype indicated. Dot plots = raw data from 3 separate embryos. Line graphs underneath = averages of these data. (G) Left. In wildtype embryos Arm accumulation varies smoothly over the segment. Middle. Loss of Wg flattens the Arm stripes. Right. Expressing Axin:GFP using the MatGAL4 driver blunts or eliminates Arm stripes. (H) The slightly lower levels of Axin expression in Mat RFP&Axin embryos have more variable effects on Arm stripes. (I-L) Over a threshold, elevating Axin levels reduces Arm accumulation in Wg stripes but does not affect interstripes. (I,K) Box and whisker plot comparing Arm accumulation levels in Wg-expressing stripes versus Arm levels in the interstripes for the indicated genotypes. n=10 pairs. Boxes extend from 25^th^ to 75^th^ percentiles, and whiskers indicate minimum to maximum values. Median=middle line of the box and mean=+. (J,L) Scatter plots showing difference in Arm accumulation between the Wg stripes and interstripes within individual embryos. Each point = a single embryo. Error bars=mean+S.D. Statistical analysis: A paired t-test was used to determine the significance between intragroup values in I and K. To assess the significance between intergroup values, an unpaired t-test was used in I and J, and an ordinary one-way ANOVA followed by Dunnett’s multiple comparisons test was applied in K and L. ns, not significant i.e. p**=** 0.05. *= p< 0.05. **= p< 0.01. ***= p< 0.001. ****=p< 0.0001. Scale bars=30µm.

We thus developed methods to quantify the effects of elevating Axin levels on two different aspects of Arm stabilization. To quantify the <u>graded effects</u> of Wg signal across the full segment (Fig. 4B”, curve), we used a digital image mask (Fig. S2A’) to remove the cortical Arm in cell-cell adherens junctions (Fig. S2A vs. A”), and then measured fluorescence levels of cytoplasmic and nuclear Arm pixel by pixel across two to three body segments (Fig S2A” box; two wildtype examples are in Fig. 4G left). In wildtype embryos, both our images and quantitative analysis revealed a smooth gradation of Arm accumulation, from peaks centered on Wg stripes to troughs in the interstripes (Fig. 4A,B,G). As a control, we examined *wg* null mutants, in which Arm levels were not elevated in any cells (Fig. 4G; each mutant was analyzed in parallel with the wildtype shown to its left). 9-fold elevation of Axin (Mat Axin) led to either complete loss of this graded stabilization of Arm in cells receiving Wg signals, or a reduction in the height of the peaks, relative to wildtype (Fig. 4C,D, quantified in G). The changes in Arm peak heights were dependent on the level of Axin:GFP expression; this was best visualized in Mat RFP&Axin embryos where the lower level Axin expression only partially flattened the Arm distribution (Fig. 4E-F. quantified in H).

To measure <u>absolute levels</u> of Arm stabilization by Wg signaling, we assessed Arm fluorescence in two groups of cells: 1-2 cell rows centered on cells expressing Wg (the Wg stripes; Fig. S2B, B’, yellow boxes) and 1-2 cell rows farthest from the Wg-expressing cells (the interstripes; Fig. S2B, white boxes). Wildtype embryos were included on the same slides as a control. We quantified both absolute Arm levels in both Wg stripes and interstripes (Fig. 4B”, black arrows, I, Table S5) and the difference in levels between these two cell types (Fig. 4B’ red arrow, J, Table S6). 9-fold overexpression of Axin (Mat Axin) substantially reduced Arm accumulation in Wg stripes, to levels similar to those normally seen in interstripes (Fig 4C,D vs. A; quantified in I-J, Tables S5-6). However, strikingly, Arm accumulation in interstripes was unaffected. The 4-fold Axin overexpression in Mat RFP&Axin embryos also reduced Wg-stabilization of Arm, but when we sorted embryos by level of Axin:GFP expression, this was less pronounced in embryos with lower levels of Axin:GFP (Fig. 4E,F vs. A, quantified in K,L, Tables S5-6).

Together, these data suggest that there is a fairly sharp threshold of Axin levels over which the destruction complex cannot be effectively inactivated by Wg signaling. However, it was also striking that elevating Axin levels did not further increase Arm destruction in cells not receiving Wg signal (Fig. 4I, Table S5), suggesting the destruction complex in those cells may already be operating at maximal efficiency.

### Levels of APC2 can be substantially elevated without significantly affecting viability or Wg-regulated cell fates

We next investigated whether Wg signaling was similarly affected by altered APC2 levels—since it is the other key component of the destruction complex, it was important to assess whether it is also rate-limiting. We used a similar approach to mis-express GFP:APC2. Using the MatGAL4 driver, we achieved a roughly 12-fold increase in levels of APC2 (Fig 2D,E; Table S1; hereafter Mat APC2). As we observed with Mat Axin, in Mat APC2 progeny GFP:APC2 levels started high and slowly decreased (Fig. 2G). Strikingly, elevating APC2 levels more than 10-fold had no effect on embryonic viability (94% viable; Fig. 3A, Table S2); in fact, these embryos could grow up to adulthood and produce viable offspring. We next examined whether elevating APC2 levels had any effect on Wg-regulated cell fate choices, as assessed by examining cuticle phenotypes. Little or no effect on embryonic patterning was seen (Fig. 3B,C Table S3), and the few denticle belt fusions observed were in hatched larvae. Finally, we examined the effect on expression of the product of the Wg target gene *engrailed*. This was also unaffected by overexpression of APC2 (Fig 3D, J, Table S4). Thus, in stark contrast to Axin, embryonic viability, cell fate choice and Wg target gene expression are not sensitive to substantially elevated levels of APC2.

### Elevating levels of APC2 strongly promotes downregulation of the destruction complex oin response to Wg signaling

As a final exploration of the effects of elevating APC2 levels, we examined Wg-regulation of Arm stability, using the same assays we employed for analyzing effects of altering Axin levels (Fig. S2). We were surprised to find APC2 overexpression led to a striking change in Arm levels and thus activity of the destruction complex. Levels of Arm in Wg-expressing cells and their immediately adjacent neighbors were sharply <u>elevated</u> (Fig. 5B-D vs. A), leading to a much <u>sharper</u> and more exaggerated pattern of differences in Arm accumulation across each segment. Quantification confirmed that while interstripe Arm levels were unchanged, Arm levels in Wg stripes were significantly higher (Fig. 5E,F, Table S5-6). The sharpened stripes and elevated Arm levels in Wg-ON cells were also apparent in our analysis of Arm levels across each segment (Fig. 5G). These data were quite surprising, as they were the exact opposite of the effects of elevating Axin levels. They suggest that increasing the APC2:Axin ratio stabilizes Arm in cells receiving Wg signaling, potentially by <u>increasing</u> the ability of Wg signaling to downregulate the destruction complex. They also suggest that further elevating Arm levels in cells already receiving Wg signals has little effect on Wnt-target gene expression or cell fate. Finally, they suggest that elevating APC2 levels does not alter destruction complex activity in cells not receiving Wg signals.

**Figure 5:**
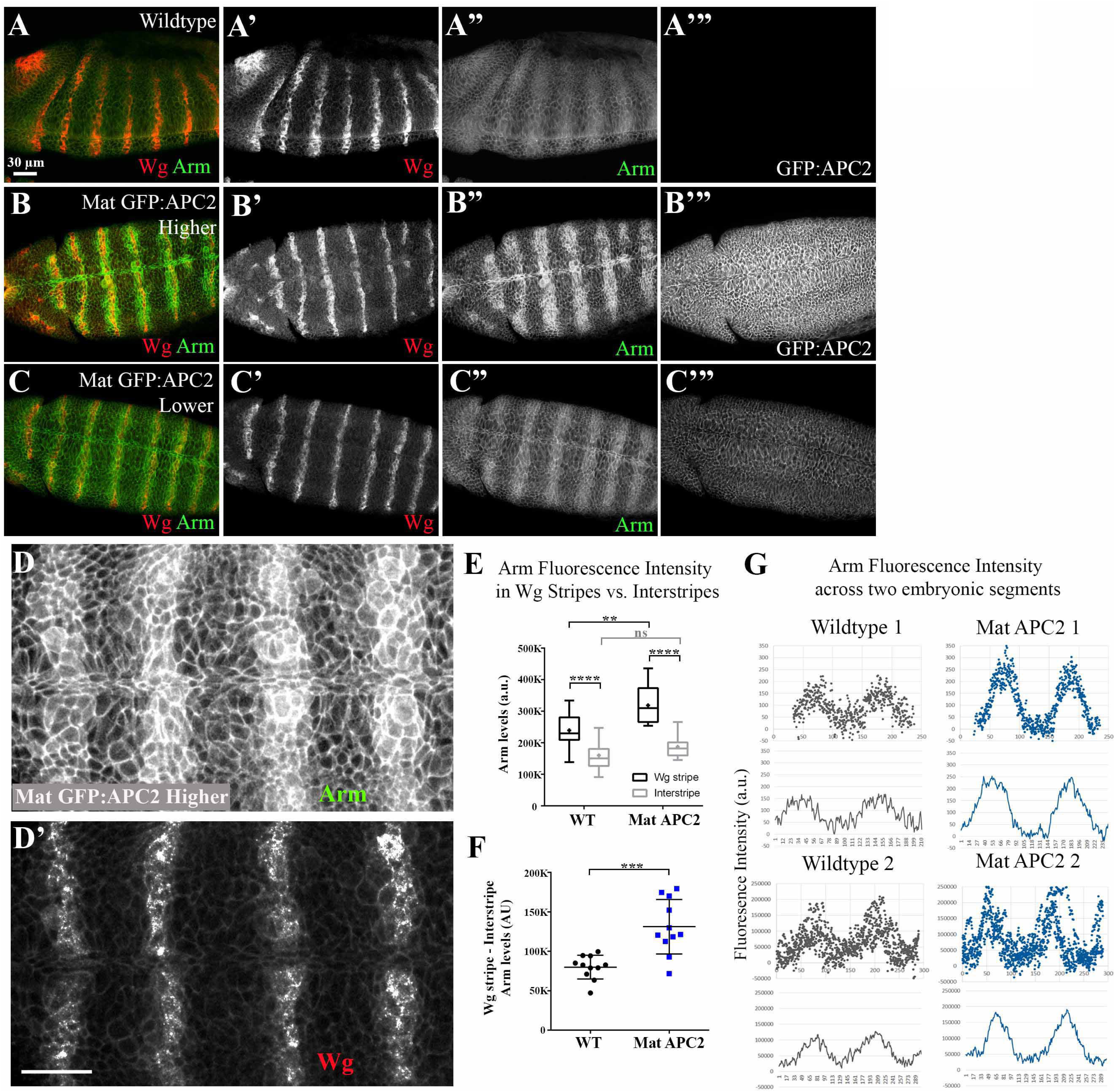
Elevating APC2 levels increases the ability of Wg signaling to turn off the destruction complex and thus increases Arm levels in those cells. (A-D) Fixed images, Stage 9 embryos. Anterior to the left. (A-C) Representative images, wildtype (A) or Mat GFP:APC2 embryos with higher (B) or lower (C) levels of GFP:APC2 expression. Elevating APC2 levels increases levels of Arm specifically in cells receiving Wg signal, (D) Close-up, embryo expressing elevated levels of GFP:APC2. The boundary of cells with elevated levels of Arm is quite sharp, and does not expand much farther than the cells adjacent to those expressing Wg. (E) Elevating APC2 levels increases Arm accumulation in Wg stripes but does not affect interstripes. Box and whisker plot (as in Fig. 4G), comparing Arm accumulation levels in Wg-expressing stripes versus Arm in the interstripes in wildtype or Mat GFP:APC2 embryos imaged on the same slide. (F) Difference in Arm accumulation between the Wg stripes and interstripes within individual embryos. Each point = a single embryo. (G) Plots of Arm accumulation pattern over 2 segments. Dot plots = raw data from 3 separate embryos. Line graphs underneath = averages of these data. Elevating APC2 levels exaggerates and sharpens the Arm stripes. Statistical analysis: a paired t-test was used to determine the significance between intragroup values in E, and an unpaired t-test was used to determine the significance between intergroup values in E and F. ns, not significant i.e. p= 0.05. **=p< 0.01. ***= p< 0.001. ****=p< 0.0001.

### Simultaneously elevating levels of <u>both</u> APC2 and Axin inhibits Wg signaling more than elevating levels of Axin alone

Thus, elevating Axin levels or elevating APC2 levels had <u>opposite</u> effects on the ability of Wg signaling to reduce destruction complex function. To explore this further, we sought to vary the levels of both proteins simultaneously, and also to vary the ratios of their expression levels. We began by expressing both Axin:GFP and GFP:APC2 simultaneously, in the progeny of GFP:APC2/MatGal4; Axin:GFP/Mat Gal4 females crossed to GFP:APC2; Axin:GFP males (hereafter, Mat APC2 & Axin). The progeny of this cross will differ in their zygotic genotypes and thus in the relative levels of Axin:GFP and GFP:APC2 (Fig. 6A). We first examined the average levels of overexpression in embryos of all four genotypes combined. Immunoblotting revealed that progeny of this cross, <u>on average</u>, accumulate Axin:GFP at levels 4-fold above endogenous Axin (Fig. 2H-J, Table S1) similar to Mat RFP&Axin (which also contains two UAS transgenes), and accumulate GFP:APC2 at ∼20x endogenous levels (Fig. 2I, Table S1). However, embryonic lethality of embryos overexpressing both Axin:GFP and GFP:APC2 was substantially higher than that of Mat RFP&Axin embryos (63% versus 32% lethal; Fig. 6C vs. Fig. 3A; Table S2), despite similar average levels of Axin:GFP accumulation (Fig. 2H-J, Table S1). In parallel, cell fates were shifted more towards the *wg* null phenotype (Fig. 6D, Table S3) than was seen in Mat RFP&Axin embryos (Fig. 3B, Table S3). Therefore, co-expression of APC2 and Axin inhibits Wg signaling to a greater extent than expression of Axin alone or APC2 alone, despite similar average levels of Axin:GFP and GFP:APC2 accumulation (Fig. 2H-J, Table S1). These data are consistent with the hypothesis that co-expressing Axin and APC2 enhances the resistance of the destruction complex to inactivation by Wg signal.

**Figure 6:**
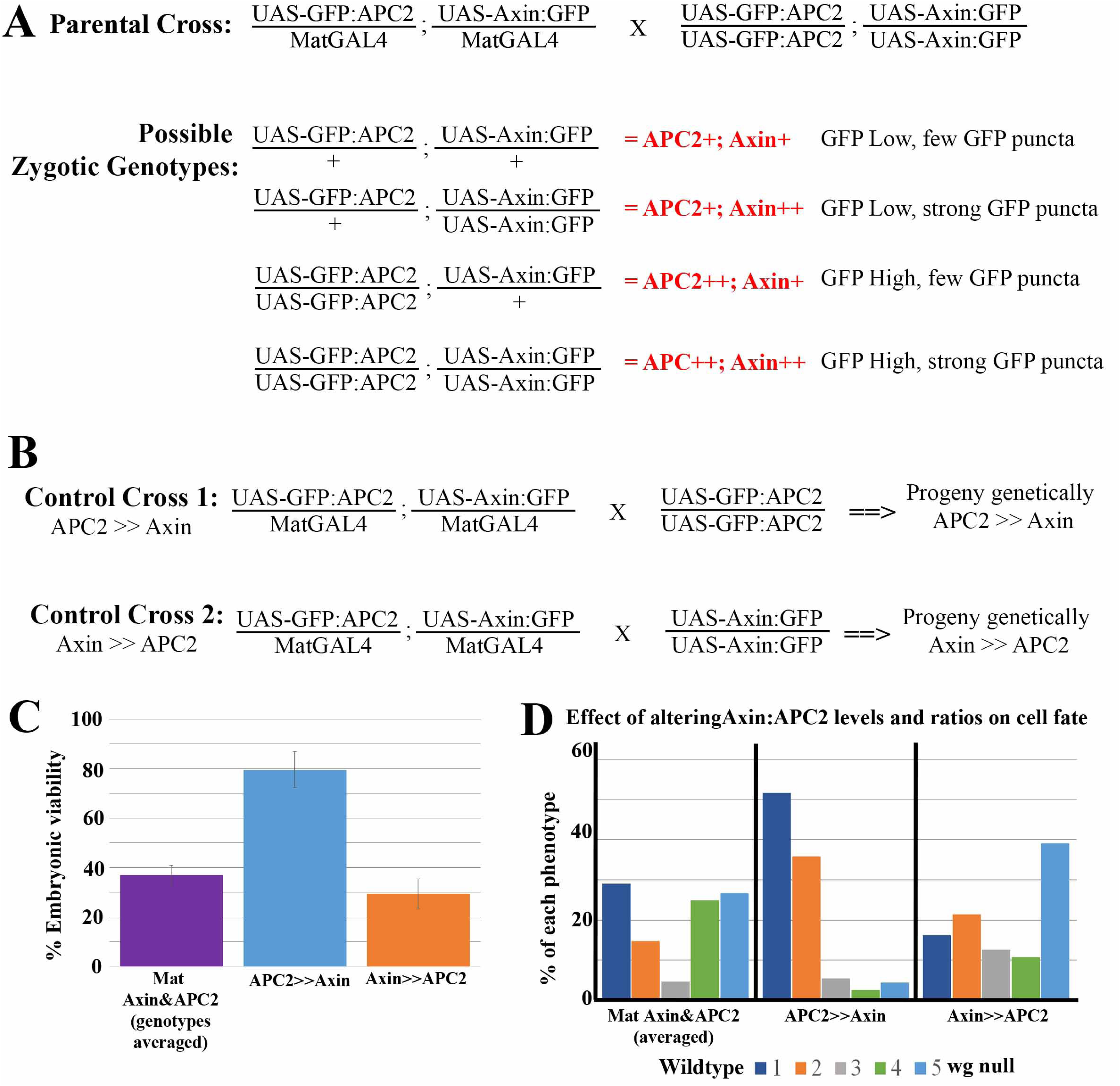
The relative ratios of APC2 to Axin levels determine effects on embryonic viability and Wg-regulated cell fates. (A) Cross used to generate embryos expressing different ratios of APC2 and Axin, with the four categories of progeny, their relative levels of Axin and APC2 overexpression, and the criteria used to identify them. (B) Control crosses used to assess how different ratios of Axin and APC2 overexpression differentially affect embryonic lethality and Wg-regulated cell fate choice. (C) Embryonic viability of different genotypes with differentially altered APC2:Axin ratios. (D) Quantification of the effects of elevating APC2 and Axin on cell fate, as assessed by cuticle pattern. Representative cuticles are in Fig. 3C.

We suspected that these averages hid differences in outcome among the four different genotypes present among the embryonic progeny (Fig. 6A), which would express different ratios of APC2 and Axin. To determine which genotypes exhibited embryonic lethality and defects in Wg-regulated cell fates, we set up two additional crosses, in which the relative zygotic expression of Axin and APC2 differed (Fig. 6B): 1) APC2>>Axin = average zygotic dose of GFP:APC2 <u>higher</u> than that of Axin:GFP, and 2) Axin>>APC2 = average zygotic dose of GFP:APC2 <u>lower</u> than that of Axin:GFP. These two crosses had strikingly different results. APC2>>Axin progeny had only 24% embryonic lethality and Wg-regulated cell fates were only mildly affected (Fig. 6C,D, Table S2-3), while Axin>>APC2 progeny had 78% embryonic lethality and had very strong effects on Wg-regulated fates, with 44% having a *wg* null phenotype (Fig. 6C,D, Table S2-3). Thus, while APC2 overexpression alone does not affect cell fates, elevating levels of both APC2 and Axin levels inhibits Wg signaling to a greater degree than elevating levels of Axin alone, suggesting that both the <u>total levels</u> and the <u>relative ratios</u> of Axin and APC2 have effects.

### The relative ratio of APC2:Axin levels determines the effectiveness of Arm destruction

These data made strong predictions about how different relative levels of Axin and APC2 would affect Arm destruction. While we could not directly determine genotypes of fixed and stained embryos, we developed a method to infer genotypes from levels and localization of GFP-tagged proteins. Since total protein levels of GFP:APC2 were, on average, overall higher than those of Axin (Figure 2I, Fig. S3A vs. B), we first separated embryos into two categories, by directly quantifying total GFP expression levels by immunofluorescence, and then using low versus high GFP levels as a surrogate for zygotically UAS-GFP:APC2/+ versus zygotically UAS-GFP:APC2/UAS-GFP:APC2 embryos (e.g., Fig 7A ‘“, B’” vs. C’”,D’”). To further subdivide the embryos, we made use of the assembly of Axin:GFP into cytoplasmic puncta [13]. If we could easily visualize cytoplasmic puncta (Fig. 7B’”,D’” insets), we categorized these embryos as zygotically UAS-Axin:GFP/UAS-Axin:GFP rather than zygotically UAS-Axin:GFP/+. This produced four presumptive genotypes with different degrees of overexpression of Axin and APC2 (Fig. 6A):

1. APC2+Axin+. Presumptive zygotic genotype = UAS-GFP:APC2/+; UAS-Axin:GFP/+,
2. APC2+Axin++. Presumptive zygotic genotype = UAS-GFP:APC2/ +; UAS-Axin:GFP/UAS-Axin:GFP.
3. APC2++Axin+. Presumptive zygotic genotype = UAS-GFP:APC2/UAS-GFP:APC2; UAS-Axin:GFP/+
4. APC2++ Axin++. Presumptive zygotic genotype = UAS-GFP:APC2/UAS-GFP:APC2; UAS-Axin:GFP/UAS-Axin:GFP.

**Figure 7:**
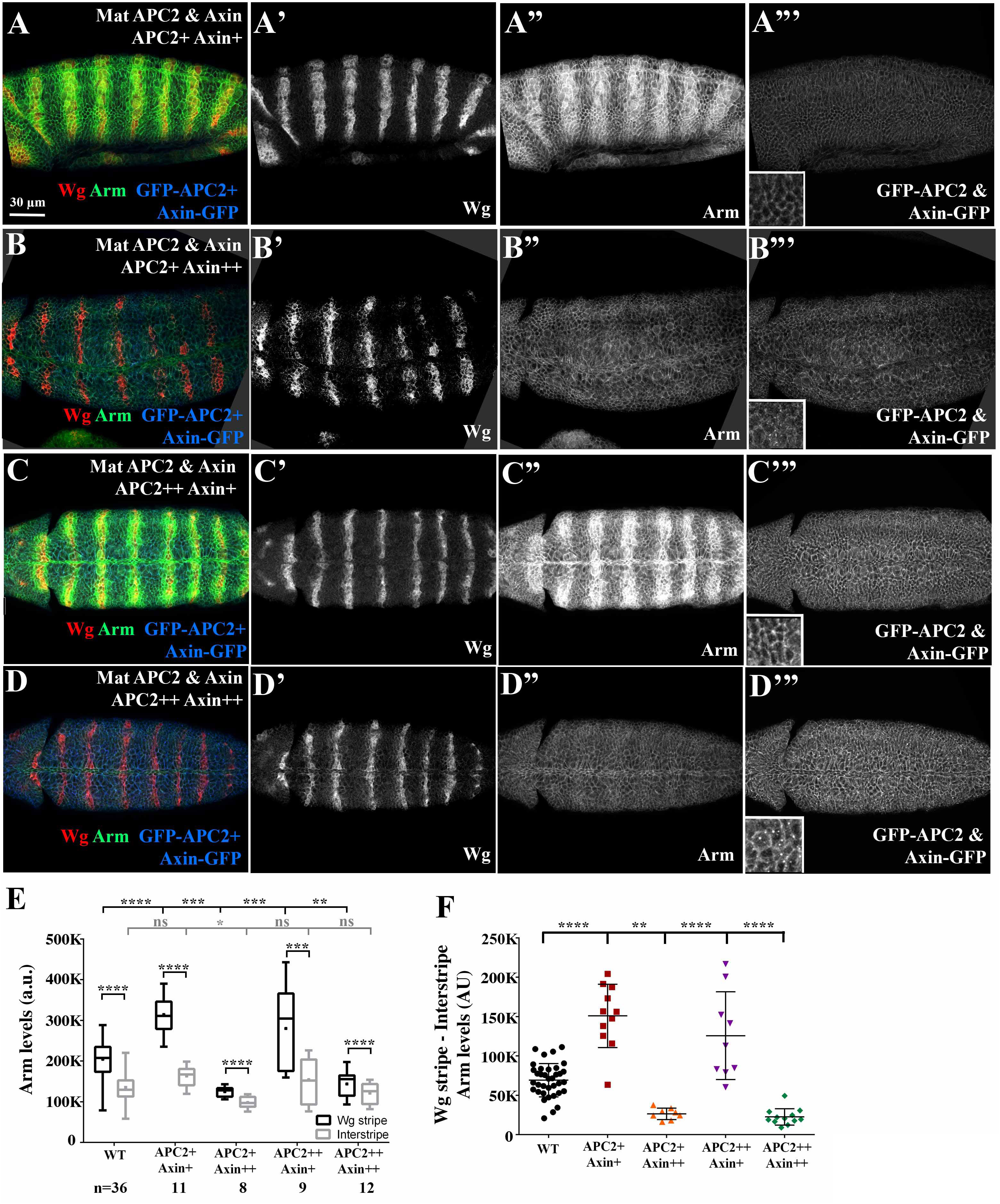
The relative ratios of APC2 to Axin levels determine effects on Arm destruction. (A-D) Fixed, stage 9 embryos. Anterior to the left. Representative images of the four different categories of the Mat APC2 & Axin phenotypes. Images were taken under the same microscope conditions. Insets are close-ups. See Fig. 6A for key to identifying presumptive genotype. (A,C) Both genotypes in which GFP:APC2 elevation exceeds that of Axin:GFP have elevated Arm accumulation in Wg stripes. (B,D) Both genotypes with the highest levels of Axin:GFP have reduced Arm accumulation in Wg stripes. (E) Effects on Arm accumulation in Wg stripes or interstripes in embryos with different ratios of Axin and APC2 accumulation. Box and whisker plot (as in Fig. 4G), Arm accumulation in Wg-expressing stripes versus Arm in the interstripes. Wildtype and different presumptive genotypes of Mat APC2&Axin embryos imaged on the same slide. (F) Difference in Arm accumulation between Wg stripes and interstripes within individual embryos. Statistical analysis, a paired t-test was used to determine the significance between intragroup values in G, and an ordinary one-way ANOVA followed by Dunnett’s multiple comparisons test was applied between intergroup values in G and H. ns, not significant i.e. p**=** 0.05. **= p< 0.01. ***= p< 0.001. ****=p< 0.0001.

We then analyzed Arm accumulation in these four embryo categories, using the quantitative tools described above to assess absolute Arm levels in Wg stripes and interstripes relative to wildtype controls. To our surprise, despite the four presumptive genotypes, the embryos divided into two phenotypic categories with regard to Arm accumulation. In embryos of the two genotypes that overexpressed Axin at the highest levels (APC2+Axin++, (Fig. 7B); and APC2++Axin++, (Fig 7D)), Arm levels were <u>strongly reduced</u> in the Wg stripes (Fig. 7B,D: quantified in Fig 7E,F; Tables S5-6). Thus, they resembled embryos overexpressing only Axin (Fig. 4C,D,I,J). In contrast, the two genotypes that overexpressed APC2 but had lower levels of Axin elevation (APC2+Axin+ (Fig. 7A) and APC2++Axin+ (Fig. 7C)), Arm levels were <u>strongly elevated</u> in the Wg stripes (Fig. 7A,C: quantified in Fig 7E,F, Tables S5-6). Thus, they resembled embryos overexpressing APC2 alone (Fig. 5B, C). Combined with the phenotypic data above, these data suggest that the ratio of APC2 to Axin plays a very important role in determining sensitivity of the destruction complex to being inactivated by Wg signaling. We next explored the effects of Wg signaling and different levels of Axin and APC2 on subcellular localization of Axin.

### Axin assembles into cytoplasmic multiprotein destruction complexes, and Wnt/Wg signaling leads to their membrane-recruitment and elevates levels of cytoplasmic Axin

One major question still debated in the Wnt field is what happens to the destruction complex after Wnt stimulation. Wnt signaling leads to Axin recruitment to the transmembrane receptor LRP5/6 [18]. Work in both cultured human cells and *Drosophila* embryos suggest that both core components of the destruction complex, APC and Axin, can be recruited to the membrane after Wnt stimulation [8,13]. However, three studies of the resulting effects of Wg signaling on Axin levels and localization in the *Drosophila* embryonic epidermis yielded to three distinct conclusions: 1) Wg signaling destabilizes Axin [21], 2) Wg signaling initially stabilizes Axin [15], or 3) Wg signaling leads to membrane recruitment of Axin [13].

We thus revisited the issue, taking advantage of our ability to express Axin:GFP at known levels and below the threshold at which it inhibits Wg signaling. We first looked at stage 9 Mat RFP&Axin embryos, which express Axin at 4-fold endogenous levels. To avoid issues with antibody accessibility to Axin assembled into large multiprotein complexes versus protein diffuse in the cytoplasm, an issue we encountered in cultured colorectal cancer cells [5], we directly visualized Axin:GFP by its fluorescence. In cultured colorectal cancer cells, Axin self-assembles into multiprotein “puncta” and can recruit APC into these structures [33]. We hypothesize these puncta are larger versions of the normal multiprotein destruction complex [5,30]. Similar puncta were clearly visible in *Drosophila* embryos when Axin was significantly overexpressed [13], at levels that inhibit Wg signaling.

We thus examined Axin:GFP localization in embryos expressing it at levels that do not substantially disrupt Wg-regulated cell fates (Mat RFP&Axin = 4-fold elevated). *wg* mRNA expression initiates at the blastoderm stage. As germband extension starts, Wg protein is just beginning to accumulate in stripes (Fig. 8A,B; [31]). We found that most cells had small puncta of Axin:GFP, both membrane-proximal and cytoplasmic. In some cells near to those initiating Wg expression, Axin:GFP containing puncta were beginning to be enriched at the cortex (Fig. 8B arrows). In contrast, at stage 9, when Wg signaling begins to regulate Arm levels and shape cell fate, we observed a prominent difference in Axin:GFP localization in cells receiving or not receiving Wg signal (Fig. 8C,D). In cells far from the source of Wg, virtually all of the Axin:GFP was assembled into bright cytoplasmic puncta, with very little in the cytoplasm (Fig. 8D, E yellow arrows). In contrast, in cells receiving Wg signal, Axin:GFP assembled into smaller membrane-associated puncta, and significant levels of Axin:GFP were seen in the cytoplasm (Fig. 8D, E magenta arrows). A similar pattern was observed using GAL4 drivers that led to higher levels of Axin:GFP (*act5c*-GAL4 =Mat/Zyg Axin or MatGAL4 without RFP = Mat Axin; data not shown). This resembled the pattern observed by Cliffe et al. (2003) using a strong GAL4 driver [13]. During stage 10, when the Wg expressing stripes become interrupted, with separate midline and lateral stripes (Fig. 8G, brackets), the pattern of Axin:GFP localization became more complex in parallel. Differences in intracellular localization remained between cells near those expressing Wg (Fig. 8G, magenta arrows) and those farther away (yellow arrows).

**Figure 8:**
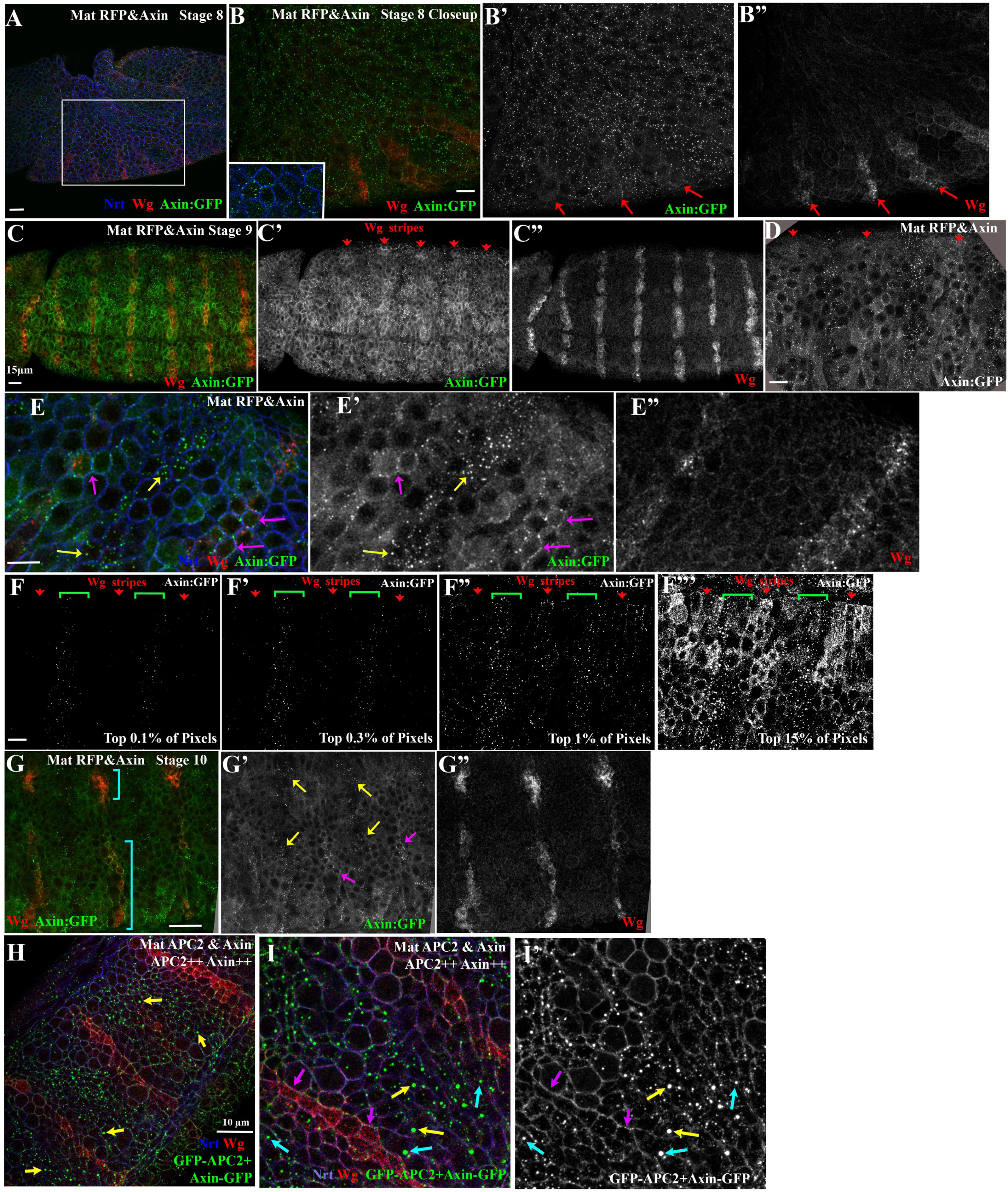
Axin assembles into cytoplasmic multiprotein destruction complexes, and Wg signaling leads to their membrane-recruitment and elevating levels of cytoplasmic Axin. (A,B) Fixed stage 8 Mat RFP&Axin embryo. Axin:GFP accumulates in puncta in all cells. Anterior to the left. Inset= close-up of B. Red arrows = start of Wg stripes. (C-E) Fixed images, stage 9 Mat RFP&Axin embryos. Anterior to the left. Red arrows= Wg expressing cells. (D,E) Close-ups, embryo in C showing Axin:GFP localization change in response to Wg. Yellow arrows = cytoplasmic puncta. Magenta arrows = membrane-associated Axin:GFP puncta. (F-F”’) Image thresholding to determine the relative brightness of different pools of Axin:GFP. (F,F’) The brightest Axin:GFP pixels are in the cytoplasmic puncta in the interstripe cells (brackets). (F”) The next brightest pixels are in membrane–associated puncta in the Wg stripe cells (arrows). (F’”) Diffuse cytoplasmic staining is higher in Wg-stripe cells (arrows) than in interstripes (brackets). (G) Fixed stage 10 embryo. As the Wg stripe separates into medial and lateral domains (brackets), Axin:GFP continues to exhibit differential localization near or distant from Wg-expressing cells. Anterior to the left. Yellow arrows = cytoplasmic puncta. Magenta arrows = membrane-associated Axin:GFP puncta. (H,I) Mat APC2&Axin. Presumptive APC2++ Axin++ embryos. Simultaneously highly elevating levels of <u>both</u> APC2 and Axin enhances resistance of the destruction complex to be turned off by Wg signaling. Yellow arrow = very bright cytoplasmic puncta. Cyan arrows = bright puncta found near Wg-positive cells. Magenta= membrane-associated puncta in Wg expressing cells.

To quantitatively assess levels of Axin:GFP in different subcellular structures, we thresholded our images to different degrees, assessing which structures were brightest and thus likely contained the highest density of Axin:GFP proteins. The results were quite striking. The brightest 0.1% of pixels and most of the brightest 0.3% of pixels, which represent the highest levels of Axin:GFP accumulation, were located in the cytoplasmic puncta in Wg-OFF cells (Fig. 8F, F’). When we lowered the threshold intensity to visualize the brightest 1% of pixels, the next structures to appear were the membrane-associated puncta in Wg-ON cells (Fig. 8F”). It was only when we visualized the brightest 15% of the pixels that the relatively high levels of diffuse cytoplasmic Axin:GFP in the Wg-ON cells were revealed (Fig. 8F”’). This contrasted with the much lower cytoplasmic levels of Axin:GFP in Wg-OFF cells.

We next sought to reconcile our observations with recent publications, whose data suggested that the primary effect of Wg signaling was to stabilize Axin in both the cytoplasm and at the membrane [15,22]. These studies used an antibody to an epitope to visualize epitope-tagged Axin. We therefore used a GFP-antibody to visualize Axin:GFP expression (Fig S4A-E). Intriguingly, the bright Axin cytoplasmic puncta in the interstripe regions were much less apparent (e.g, Fig. 8C’ vs. Fig S4A’ or C’)—thus use of an antibody emphasized the stronger cytoplasmic signal in Wg-ON cells, reproducing the earlier observations. This suggested that direct visualizing Axin:GFP provides a more complete picture of the effects of Wg signaling on Axin localization and levels.

Together, these data suggest in the absence of Wg signals, Axin self-assembles into large cytoplasmic multiprotein destruction complexes and diffuse cytoplasmic levels of Axin diminish—earlier work suggested that the Axin puncta also contain APC2 [13,34]. In contrast, in cells receiving Wg signal, Axin puncta are recruited to the plasma membrane, these puncta diminish in intensity, and the cytoplasmic pool is correspondingly increased.

### Simultaneously elevating Axin and APC2 enhances puncta assembly and makes puncta resistant to Wg signaling

Our data above suggest that co-expressing Axin and APC2 could, if the ratios were right, lead to synergistic inhibition of Wnt signaling. We thus examined how altering elevating levels of both Axin and APC2 altered assembly and localization of the destruction complex. GFP:APC2 expressed alone was primarily cortical (Fig. 5B,C), as we observed for endogenous APC2 [23]. In embryos expressing both GFP:APC2 and Axin:GFP at strongly elevated levels (APC2++Axin++ embryos; Fig. 8H,I), we observed two notable differences from what we observed when each was expressed alone. First, the cytoplasmic puncta in Wg-OFF cells were brighter (Fig. 8H,I yellow arrows), consistent with the idea that APC2 recruitment into puncta may stimulate or stabilize Axin multimerization, as we observed in cultured SW480 cells [5]. Second, the region occupied by bright cytoplasmic puncta became much broader, expanding right up to the Wg-expressing cells (Fig. 8I, magenta arrows), and the region with membrane-associated puncta became narrower, now restricted largely to the Wg-expressing cell alone (Fig. 8I, blue arrows). Together with the phenotypic data above, these data suggest that if Axin levels *are limiting relative to those of APC2*, elevating APC2 levels makes the destruction complex <u>more susceptible</u> to being turned down by Wg signaling. In contrast, if Axin levels *are not limiting relative to those of APC2,* then elevating APC2 levels makes the destruction complex <u>less susceptible</u> to being turned down by Wg signaling; our data are also consistent with the idea that this occurs by stabilizing destruction complex assembly to the effects of Wg signaling.

### Each destruction complex punctum includes tens to hundreds of APC2 or Axin proteins

Data from both cultured cells and *Drosophila* suggest that the ability of Axin and APC to polymerize into a large multimeric complex is critical for targeting βcat for destruction. Overexpressing *Drosophila* Axin in colorectal cancer cells leads to assembly into large “puncta”, which we hypothesize are enlarged versions of the normal destruction complex. APC2 is recruited into these. Previous work from our lab using super resolution microscopy allowed us to begin to look inside these puncta, revealing that APC2 and Axin formed intertwining filaments [5]. To fully understand destruction complex assembly and function, one key parameter is to obtain an estimate of the number of proteins assembled into an active destruction complex. Previous work provided no insights into this issue, either with respect to the large puncta observed after overexpression in colorectal cancer cells, or the presumably smaller complexes produced when Axin and APC2 are expressed at endogenous levels.

To estimate the number of APC2 or Axin molecules within an active destruction complex, we adapted a fluorescence comparison technique developed to quantify numbers of GFP-tagged proteins in multimeric complexes [35,36]. They utilized macromolecular structures containing a known number of GFP molecules as standards (e.g., purified eGFP= 2 molecules and a virus-like particle=120 molecules), and then from these developed methods to define the number of proteins in yeast multiprotein complexes where molecule number had not been previously defined. We used 2 yeast strains from this study as standards (Fig. 9A): one expressing Ndc80:GFP (calculated to have 306 molecules) and the other expressing Mif2:GFP (calculated to have 58 molecules) [35]. Since we thought it likely that destruction complexes did not have a fixed size, our goal was to get an order of magnitude estimate of the number of proteins in each destruction complex punctum.

**Figure 9:**
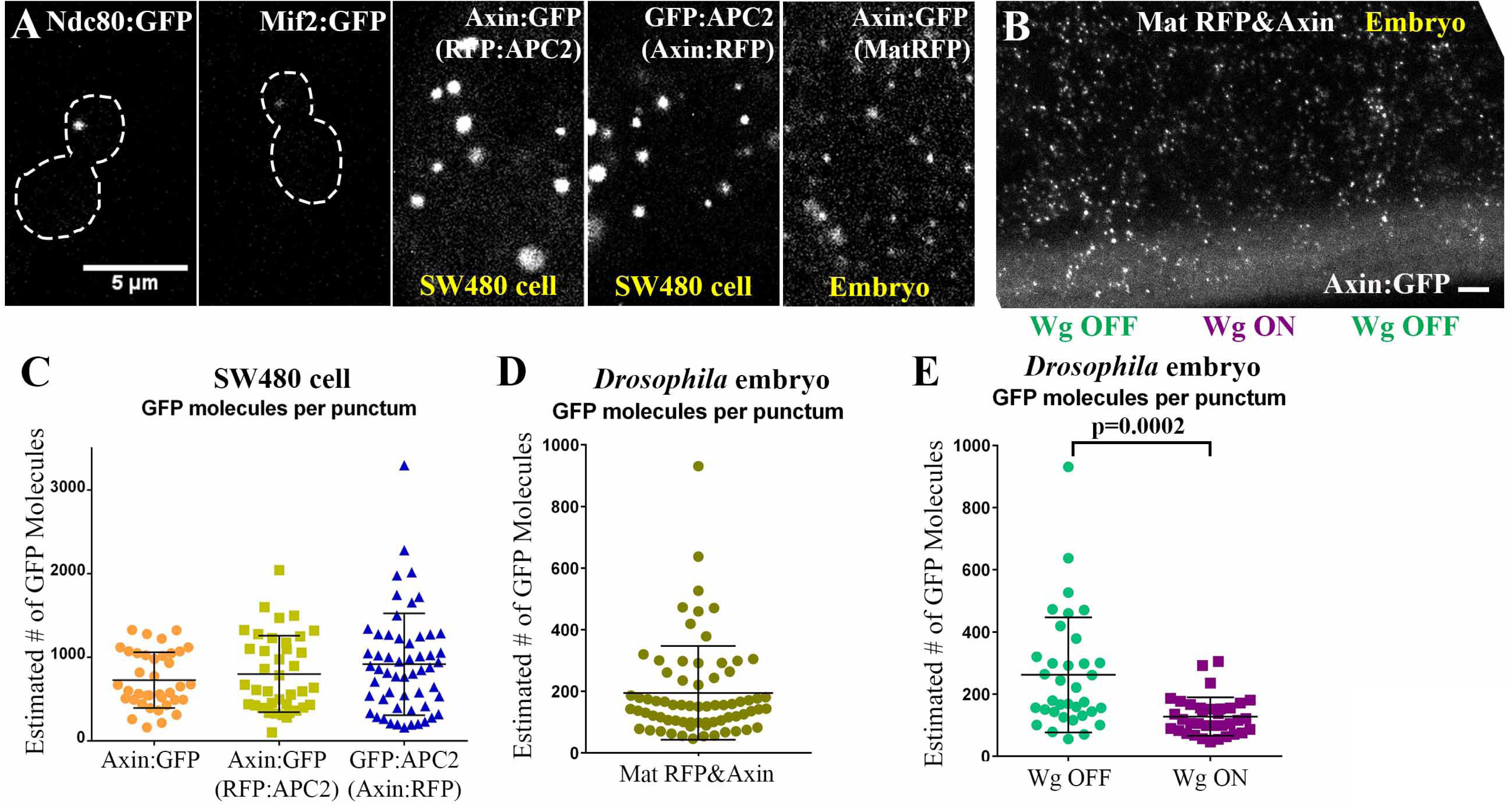
The destruction complex contains thousands of APC2 or Axin molecules after over-expression in SW480 cells, and 10-100s of Axin molecules in vivo in embryos. (A) Representative images of live samples used for fluorescence comparisons to calculate GFP molecule numbers. Each panel is scaled to the same size and brightness. Ndc80:GFP assembles into a structure containing 306 GFP molecules while Mif2:GFP assembles into a structure containing 58 GFP molecules. (B) Pattern of Axin:GFP accumulation and localization in a live embryo. Comparison to our fixed samples allowed identification of regions receiving Wg signal(dimmer puncta) or not receiving Wg signal (brighter puncta) (C-E) Estimated number of GFP molecules per punctum. Each dot represents an individual punctum analyzed. Means and standard deviation are in Table S7. (C) GFP Molecule counts from SW480 colorectal cancer cells expressing Axin:GFP alone, Axin:GFP plus RFP:APC2, or GFP:APC2 in addition to Axin:RFP. (D-E) GFP molecule counts *in vivo* from stage 9 embryos expressing RFP and Axin:GFP under the control of Mat-GAL4 (Mat RFP&Axin). (E) Quantification of puncta GFP molecule counts from D, after being separated into those in presumptive regions receiving or not receiving Wg signals (as in B). Statistical analysis via an unpaired t-test test.

We first examined GFP-tagged *Drosophila* Axin over-expressed in SW480 cells. Axin uses its DIX domain to polymerize, forming cytoplasmic puncta in a large range of sizes and brightnesses [5]. We compared living yeast and Axin-expressing SW480 cells in parallel (Fig. 9A), using identical imaging conditions. Puncta size in these cells varies over several orders of magnitude [5], and thus the brightest puncta in each cell were too bright to quantify using our yeast standards. We determined brightness of individual puncta and used the two yeast standards to estimate relative brightness and thus relative molecule number. This allowed us to obtain order-of magnitude estimates of the number of Axin molecules per punctum. In the set we analyzed, the number of Axin:GFP molecules per punctum ranged from 163-1327 (mean ∼700; Fig. 9C; Table S7). When APC2 is expressed along with Axin in SW480 cells, it is recruited into the Axin puncta [30]. We thus also examined SW480 cells coexpressing both to get order of magnitude comparisons of the number of Axin or APC2 molecules in puncta. In cells co-transfected for GFP:Axin and RFP:APC2, the number of GFP:Axin molecules ranged from 104-2041 (Fig. 9C; Table S7), while in cells transfected with a GFP:APC2 and Axin:RFP, the number of GFP:APC2 molecules per punctum ranged from 162-3297 (Fig. 9C; Table S7), suggesting puncta contain roughly comparable numbers of both proteins. Because the brightest puncta were outside the range quantifiable using our yeast standards, these numbers provide a lower limit for molecule number in the largest puncta. These data suggest that when over-expressed in SW480 cells, APC2 and Axin can assemble into destruction complexes containing at least 100s to 1000s of each protein, and within the complex are likely to be present at the same order of magnitude in molecule number.

While this offered insights into the assembly ability of these two proteins, it involved very significant overexpression in an *APC* mutant colorectal cancer cell line. To assess molecule numbers in an active destruction complex in a natural context and at more normal levels of expression, we turned to live *Drosophila* embryos from the Mat RFP&Axin line. These embryos express Axin:GFP at 4 times above endogenous levels and more than 60% of these embryos are viable with no or subtle defects in Wg-regulated cell fates (Fig. 2A,B 3A). We imaged Mat RFP&Axin embryos live, in parallel with yeast expressing each of our two protein number standards (Fig. 9A). Fluorescence comparison revealed that the Axin:GFP puncta range from 46-931 molecules of Axin per punctum (at stage 9; average ∼200; Fig. 9D, Table S7). As noted above, subcellular localization and apparent brightness of Axin:GFP puncta changed in response to Wg signaling, with the brightest puncta in the cytoplasm of Wg-OFF cells and dimmer, membrane-bound puncta in Wg-ON cells. This difference across the segment was apparent in our live Mat RFP&Axin flies (Figure 8B). We used these criteria to separate the puncta into those in Wg-ON versus Wg-OFF cells. There was a significant difference between the numbers of Axin:GFP molecules in puncta found in Wg OFF (average ∼260 molecules) versus Wg ON regions (average ∼130 molecules; Fig. 9E; Table S7), although the distributions overlapped. These data provide the first insight into the scale of macromolecular assembly in an endogenous destruction complex, suggesting each contains 10s to 100s of Axin molecules. They also support the idea that the number of Axin molecules per destruction complex decreases in response to Wg signaling.

## Discussion

Wnt signaling plays key roles in cell fate choice and stem cell homeostasis in normal development, and mutational activation underlies colorectal and other cancers. The key regulated step in signaling is regulation of the stability of the Wnt effector ßcat by the multiprotein destruction complex. Despite the intense interest in this pathway, the mechanisms by which Wnt signaling regulates destruction complex activity remains a key question in the Wnt field. Our data provide new insights into this in several ways.

### In vivo levels of APC and Axin are similar rather than orders of magnitude different

Pioneering work *Xenopus* egg extracts defined key parameters underlying the biochemical action of the destruction complex, by assembling and measuring destruction complex activity. In these studies, they lacked reagents to directly measure protein levels of all of the components, and thus used addition of purified Axin to estimate its relative levels. These data suggested Axin is present at levels much lower than the other components of the destruction complex, with an APC:Axin ratio ∼5000:1 [10,11]. Most mathematical and other models in the field use these data as an underlying premise, and thus they have shaped thinking about Wnt signaling for more than a decade.

However, recent work in cultured mammalian cells had begun to cast doubt on the universality of this ratio—in some cell lines APC levels were slightly higher (<2-fold) than Axin while in others Axin was actually present at higher levels than APC [12]. We thus decided to use a well-characterized model where the consequences of Wnt signaling are well known: the *Drosophila* epidermis during mid-embryogenesis, when cell fate is tightly regulated by Wg signaling. Using both RNAseq and direct comparisons of protein levels, we found, in contrast to *Xenopus* oocyte extracts, that the ratio of APC:Axin is much more similar. Our protein data suggest this ratio is 5:1. In fact, this may overestimate the available level of APC2, the primary APC family member at this time. APC proteins also have distinct cytoskeletal roles [37], including at times just prior to the time we examined [38], and thus the pool of APC2 available for Wg signaling may be even lower. While it is possible that the discrepancies in our results involve the species used (*Xenopus* vs. *Drosophila*), our data and the mammalian cultured cell data suggest that the difference may be in comparing tissues where Wnt signaling is active, versus those, like *Xenopus* egg extracts, in which Wnt signaling is not yet active. Thus, future mathematical modeling of Wnt signaling should include states in which APC and Axin are present at similar levels.

### In the absence of Wg signaling, Axin assembles into large cytoplasmic multiprotein complexes that each contain tens to hundreds of Axin proteins and collectively contain much of the Axin in the cell

Previous work provided conflicting results on Axin localization in the absence and presence of Wg signaling. In interstripe cells, which receive little or no Wg signal, the destruction complex is presumably in its maximally active state. Cliffe et al. (2003) suggested that Axin and APC2 co-localize in cytoplasmic puncta in these cells [13], while others did not see any notable subcellular localization of Axin in Wg-off cells [15,21]. To address this, we examined Axin:GFP localization in embryos expressing Axin below the threshold for detrimental effects (Mat RFP&Axin; 4x endogenous levels). Our data confirm and extend the work of Cliffe et al. [13] In Wg-OFF interstripe cells, Axin assembled into large cytoplasmic puncta, presumably driven by DIX-domain mediated Axin polymerization. In these cells, levels of cytoplasmic Axin were relatively low, suggesting that much of the Axin self-assembles into puncta. Earlier work suggests that these puncta also contain APC2 [13], and APC2’s ability to multimerize may also be relevant [5,6]. Our data are consistent with the idea that puncta size is regulated by APC2, since in embryos with similar levels of Axin but elevated levels of APC2, puncta size increased, as assessed by brightness. This would be consistent with our work in cultured mammalian cells, which suggested one key role of APC family proteins in promoting destruction complex function is to stabilize Axin multimerization [5]. Our molecular counting experiments also provided us with the first opportunity to assess the number of molecules in the multiprotein destruction complex. These data suggest active destruction complexes contain tens to low hundreds of Axin proteins, thus helping explain the critical role of the Axin DIX domain [3,16], which mediates Axin polymerization [4,39]. Our recent work to engineer the minimal Wnt regulatory machine confirmed that both Axin’s DIX domain and APC2’s Arm repeats, implicated in polymerization, are among the domains most critical for destruction complex function [40].

### Wg signaling triggers membrane recruitment of Axin and may destabilize destruction complex assembly

One key and controversial question in the field involves the mechanism(s) by which Wg signaling turns down the destruction complex. Different studies in cultured mammalian cells and *Drosophila* (see Introduction) led to quite different conclusions, ranging from total disassembly of the destruction complex to inactivation of an intact complex to stabilization of Axin. Our new tools allowed us to examine Axin localization directly using a GFP-tagged protein expressed at near endogenous levels, in a tissue where we can examine cells before the onset of Wg signaling, as well as in side-by-side cells experiencing high or low levels of signaling. Our data suggest that in this tissue, Wg signaling leads to membrane recruitment of the destruction complex and are consistent with the idea that it destabilizes assembly, increasing the pool of cytoplasmic Axin. Before Wg signaling is initiated, Axin:GFP was in cytoplasmic puncta in all cells. However once Wg signaling initiated, Axin:GFP localization differed between cells. In cells not receiving Wg, most of the Axin is assembled into large cytoplasmic puncta, leaving relatively low levels diffuse in the cytoplasm. However, in cells receiving Wg signal, the remaining Axin puncta were recruited to the membrane. Our molecular counting and image thresholding experiments suggest these dimmer membrane-proximal puncta contain fewer Axin molecules. Our image thresholding experiments further suggest that in Wg-receiving cells, diffuse cytoplasmic levels of Axin are elevated relative to Wg-OFF cells. These data support and extend the earlier work of Cliffe et al. (2003) [13], who expressed a GFP-tagged Axin at more elevated levels. Our observation of elevated cytoplasmic levels of Axin in Wg-ON cells is consistent with earlier work [15], though we do not see clear evidence that this results from Axin stabilization. Our data further suggest that their use of antibody staining of an epitope-tagged protein rather than direct visualization emphasized these diffuse cytoplasmic pools of Axin while simultaneously de-emphasizing the larger cytoplasmic puncta, due to differential antibody accessibility. Thus, the stabilization of Axin proposed by the Ahmed lab may in fact largely involve a change in protein localization. This is consistent with the immunoblotting experiments of Cliffe et al (2003), who did not detect altered Axin levels. Based on our data, we hypothesize that in the presence of Wg signaling, Axin puncta are recruited to the membrane, presumably by binding to the activated Wg-receptor. We further hypothesize that Wg signaling either destabilizes puncta or inhibits puncta assembly, increasing the relative amount of Axin in the cytoplasmic pool.

### Above a tight threshold, elevating Axin levels render the destruction complex insensitive to inactivation by Wg signaling

Previous work revealed that sufficiently elevating Axin levels could inactivate Wnt signaling either in cultured mammalian cells [41] or in *Drosophila* embryos [13,42]. More recent work suggested that this only occurred when Axin levels were elevated over a certain threshold [14,16]. Our knowledge of absolute levels of APC2 and Axin allowed us to vary levels of each individually or together, thus varying both levels and ratios of the two proteins in the *Drosophila* embryo where effects of Wg signaling are well characterized. By assessing the effect on embryonic viability, expression of the target gene *en*, and cell fates choices, we confirmed that in order for elevated Axin levels to be detrimental in vivo, Axin levels must reach a minimum threshold. When Axin:GFP was expressed at ≤4x endogenous Axin, we observed little or no effect on any of these parameters, while at >8x endogenous Axin there was a dramatic increase in embryonic lethality, reduced En expression, and a shift towards a more *wg*-null like phenotype. Our data also provided insight into the underlying mechanism: increasing Axin above this threshold inhibited Wg signaling’s ability to turn down the destruction complex, and thus decreased Arm levels <u>specifically</u> in Wg-ON cells. Further mechanistic insights remain to be determined, but previous work suggest that a key parameter may be the levels of “active Dsh” protein, which is activated by Wg signaling and then can heteropolymerize with Axin and compete with APC [4,13]. If down-regulation involves a competition between Axin homo-multimerization and Dsh-hetero-multimerization, over the threshold Axin may saturate the available Dsh molecules and therefore inhibit its ability to inactivate the destruction complex, thus rendering a subset of destruction complexes immune to downregulation. Our data also revealed that Axin is <u>not</u> rate-limiting in Wg-OFF cells—there Arm levels were not further decreased by elevating Axin levels, suggesting that the destruction complex may already be working at maximal activity there.

### APC2 is not rate-limiting for destruction complex activity and in fact elevating its levels facilitates destruction complex inactivation

We next asked whether APC2, the second core component of the destruction complex, is also rate limiting for destruction complex activity. Expressing GFP:APC2 at>10x endogenous levels had little to no effect on embryonic lethality, En expression, or Wg-regulated cell fate choices. This might be because Axin is rate-limiting—thus additional APC2 would not trigger assembly of additional destruction complexes once it exceeded the available pool of Axin. Intriguingly, however, we observed an unexpected effect of elevated levels of APC2. In wildtype embryos, *wg*, is expressed by a single row of cells in each segment, producing a gradient of Wg. In cells that receive Wg, the destruction complex is turned down, and Arm accumulates in the cytoplasm and nucleus. This gradient of Wg normally creates a gradient of Arm accumulation. In contrast, embryos with high APC2 expression there is an essentially binary change in Arm accumulation in response to Wg signaling. Wg–expressing cells and their immediately adjacent neighbors accumulate Arm at levels ∼1.5x higher than the same cells in wildtype. However, in cells more distant from the Wg-expressing cells, Arm levels are unchanged from wildtype. These data suggest a potential positive role for APC2 in turning the destruction complex down in the presence of Wg signaling.

### Effects of altering the Axin:APC2 ratio suggest APC2 can play both positive and negative roles in Wnt regulation

These paradoxically opposite effects of elevating levels of Axin or APC2 on the ability of Wg signaling to inactivate the destruction complex were among our most surprising results. They are consistent with a model in which APC2 has dual positive and negative roles in Wnt regulation. A similar unexpected positive role of APC in Wg signaling was previously observed in the eye and wing imaginal discs [43]. Our dual over-expression of APC2 and Axin provided potential insight into the underlying mechanisms, emphasizing the importance of the relative ratios of different destruction complex proteins. In embryos with elevated expression of <u>both</u> APC2 and Axin, the destruction complex could <u>not</u> be effectively turned down by Wg signals. In fact, cytoplasmic Axin puncta became brighter, and thus likely contained more Axin:GFP proteins. These data are consistent with the known role of APC family proteins in promoting Arm/ßcat destruction, and fit well with our recent work in cultured mammalian cells, which demonstrated that APC can stabilize Axin multimerization, thus increasing destruction complex size and its effective activity [5]. Further, when both APC2 and Axin were elevated, cytoplasmic puncta were even observed in cells immediately adjacent to the Wg-expressing cells, suggesting that if Axin levels are not limiting, the destruction complex can remain active even in cells receiving relatively high levels of Wg signals. As noted above, this may involve situations where Axin levels exceed those of active Dsh.

However, if Axin was limiting, effects of APC2 elevation were quite different. Now elevated levels of APC2 allowed Wg signaling to <u>more effectively</u> turn down the destruction complex, thus elevating Arm levels. Perhaps the membrane-associated pool of APC2 brings both Axin and the destruction complex in proximity to the Wg receptors and Dsh, allowing more rapid and effective turndown of destruction complex function and assembly. Another possibility is that APC2 negatively regulates Axin levels, thus in turn regulating Arm levels, as was previously suggested based on genetic analysis in imaginal discs [43]. Our data do not rule out this possibility. Finally, this may be due in part to protection of Arm from destruction by binding to APC2 in complexes lacking Axin--the ability of APC2 to retain Arm in the cytoplasm is known to provide the ability to fine-tune Wg signaling [30,44]. To test these hypotheses, it would first be useful to directly test whether Wg signaling directly affects total Axin levels, by reducing or increasing Wg signaling in stage 9 embryos. To test the hypothesis that relative levels of Axin and Dsh are key, it would be interesting to compare total Axin and Dsh levels before and during active Wg signaling, and to examine the effect of varying Dsh levels on destruction complex assembly, localization and function. If Dsh levels are a rate-limiting factor in turning off the destruction complex, then increasing Dsh may balance the effects of elevating Axin. However, elevating Dsh levels alone may not be sufficient, if the ability of the Wg receptor to “activate” the pool of Dsh is limiting. It would also be interesting to localize Dsh, and, if as expected, it preferentially localizes to Axin puncta in Wg-ON cells, to count the number of Dsh molecules in the membrane-associated Axin puncta to determine if Dsh molecule numbers are in the same order of magnitude as Axin or APC2. We also need to explore the ratio of APC2:Axin <u>within</u> the destruction complex *in vivo*, to parallel the work after overexpression in cultured mammalian cells reported here.

### A proposed model of how Wnt signaling regulates destruction complex assembly and function

Our data suggest that the relative levels of different Wnt signaling regulators are a critical determining factor, and that modulating relative levels of different components may make cells more or less sensitive to Wnt signaling. Integrating our data with data from many labs using *Drosophila* and cultured mammalian cells, we propose the following model. During embryogenesis, cells begin with relatively similar levels of the two core scaffolds of the destruction complex (4-5x more APC2 than Axin). When Wg signaling is off, Axin self-assembles into cytoplasmic complexes of tens to hundreds of molecules, which we believe represent the functional destruction complex. In this state, most of the Axin in the cell is assembled into puncta, with little free in the cytoplasm. In these cells, we propose that there are 2 pools of APC2, a pool localized to the cortex that may mediate APC2’s cytoskeletal functions, and another that is associated with and stabilizes the assembly of the Axin puncta. This would represent the maximal-activity state of the destruction complex, and it would rapidly bind, sequester and turnover all newly synthesized Arm that is not assembled into adherens junctions. In the presence of Wg signaling, LRP5/6 is recruited to the Wg receptor Frizzled, and phosphorylation of LRP5/6 recruits Axin and Dsh to the membrane. We hypothesize that Axin membrane-recruitment involves largely intact destruction complexes. Our observations are consistent with recent data suggesting that the destruction complex is not fully disassembled in response to Wg signaling nor is Arm phosphorylation by the destruction complex completely inhibited [5,7-9]. Instead the ability of the destruction complex to target Arm for destruction is inhibited, perhaps by blocking the ability to transfer Arm to the E3 ligase. Dsh contains a DIX domain, like Axin, which allows Dsh to hetero-dimerize with Axin. We hypothesize that this Dsh:Axin interaction aids in puncta re-localization and stimulates the decrease in destruction complex size and function. Dsh may actively stimulate dissociation of destruction complexes, by competition, or it may simply reduce Axin multimerization, thus inhibiting the dynamics of destruction complex assembly. Other longer-term effects may then reinforce this initial event, including ubiquitination and destruction of Axin or inhibition of GSK3 kinase activity.

## Materials and Methods

### Fly Stocks, Embryonic Lethality, and Cuticles

All crosses were performed at 25°C. Wildtype was either *y w* or *act5c-Gal4/CyO*. The following stocks were obtained from the Bloomington Stock Center: *act5c-GAL4* (4414), *Maternal alpha tubulin GAL4* (referred to as MatGAL4; a stock carrying both of the GAL4 lines in 7062 and 7063), and UAS-Axin:GFP (7225). We also used UAS-GFP:APC2 [30] and an APC2 transgene which expresses APC2 under its endogenous promoter [45].

**Cross Abbreviations** (Female x Male):

**GFP:APC2** = *APC2 promoter-GFP:APC2; APC2*^*g10*^ *x APC2 promoter-GFP:APC2; APC2*^*g10*^

**Zyg Axin** = *UAS-Axin:GFP x act5c-GAL4/+*

**Mat RFP&Axin** = *UAS-RFP/MatGAL4; UAS-Axin:GFP/MatGAL4 x UAS-RFP; UAS-Axin:GFP*

**Mat Axin** = *UAS-Axin:GFP/MatGAL4; +/MatGAL4 x UAS-Axin:GFP*

**Mat/Zyg Axin** = *act5c-GAL4/+ x UAS-Axin:GFP*.

**Mat APC2** = *UAS-GFP:APC2/MatGAL4; +/MatGAL4 x UAS-GFP:APC2*

**Mat APC2& Axin** = *UAS-GFP:APC2/MatGAL4; UAS-Axin:GFP/MatGAL4 x UAS-GFP:APC2; UAS-Axin:GFP*

**APC2 > > Axin** = *UAS-GFP:APC2/MatGAL4; UAS-Axin:GFP/MatGAL4 x UAS-GFP:APC2*

**Axin > > APC2** = *UAS-GFP:APC2/MatGAL4; UAS-Axin:GFP/MatGAL4 x UAS-Axin:GFP*

Embryonic lethality assays and cuticle preparations were as previously described [46]. Inhibition of Wg signaling was assessed by analyzing embryonic and first instar larvae cuticles with the scoring criteria found in Fig. 3C.

### Immunostaining and Antibodies

Embryos were prepared as in [47]. Briefly, flies were allowed to lay eggs on apple juice/agar plates with yeast paste for up to 7 hours. Embryos were collected in 0.1% Triton-X in water using a paintbrush, then dechorionated for 5 minutes in 50% bleach. Embryos were fixed for 20 minutes in 1:1 heptane to 9% formaldehyde, with 8mM EGTA added to preserve GFP expression. Embryos were then devitillenized by vortexing in 1:1 heptane and methanol. Embryos were then washed in methanol followed by 0.1% Triton-X in PBS, then incubated in blocking buffer (1:1000 normal goat serum diluted in 0.1% Triton-X in PBS) for 30 minutes. Embryos were incubated in primary overnight at 4°C, washed in 0.1% Triton-X in PBS, then incubated in secondary antibody for 1 hr at room temperature. Embryos were mounted in Aqua polymount (Polyscience). Primary antibodies were: Wingless (Wg, Developmental Studies Hybridoma Bank (DSHB):4D4, 1:1000), Arm (DSHB:N27 A1, 1:75), phospho-tyrosine (pTyr, Millipore:4G10, 1:1000), En (DSHB:4D9, 1:50), GFP (Abcam:ab13970, 1:10,000), and Neurotactin (Nrt, DHSB:BP 106, 1:100).

### Assessing effects on Engrailed expression

Stage 9 embryos were stained with antibody to Engrailed and imaged on a Zeiss LSM 710 or 880 scanning confocal microscope. Images were processed using FIJI (Fiji Is Just ImageJ) as follows: maximum intensity projections 8µm thick were created and thresholded to highlight cells expressing Engrailed and eliminate background noise. Three lines parallel to the midline were drawn to intersect with bands 2 through 5 of Engrailed expressing cells relative to the head, two on either side of the embryo and one just to the left of the midline. The cells in each Engrailed band which were intersected by each line were included in our measurements. The number of cells per Engrailed stripe was then determined by averaging these three values. Embryos were blind scored. Significance was assessed using a one-way ANOVA test.

## Quantitative analysis of Arm accumulation

### Graded accumulation of Arm

To quantify effects of our manipulations on the graded accumulation of Arm across the segment, control and experimental embryos were stained in parallel for phosphotyrosine, which marks the adherens junctions, and Arm. Since 70% of Arm protein accumulates at the adherens junction [48], and we wanted to focus on the cytoplasmic and nuclear accumulation of Arm, we removed the adherens junction pool of Arm by creating a mask. First, all embryos were rotated to have the anterior on the left and the midline at 180°. Next “sum intensity” projections were created that went 8µm deep into the embryo. We next used FIJI’s trainable WEKA tool with anti-phosphotyrosine staining to create a membrane mask. This mask was overlaid and subtracted from the Arm image. Next a rectangular region of interest (ROI) was drawn (446 W x 60 H pixels, spanning approximately 3 Wg stripes and 4 cells wide) starting at the first interstripe in the thorax. A profile of the ROI was plotted. ROI profiles were adjusted for embryo length and to bring valleys to zero. See Figure S2A for a visualization this process.

### Wg Stripes versus Interstripes

To calculate the absolute levels of Arm accumulation in cells receiving or not receiving Wg signals, stage 9 embryos were collected and stained as previously described. For each genotype, we added *act5c*-GAL4/Cyo embryos to the same tube as a wildtype control, allowing immunostaining and microscopy imaging on the same slide. Control and experimental embryos were distinguished by the presence or absence of GFP-fluorescence. To calculate the level of Arm accumulation, we choose a specified boxed region (100 pixels wide x 30 pixels high) spanning the width of the Wg-expressing cells, and measured the mean gray value of Arm by FIJI (Fig. S2B). Three Wg stripe regions from parasegments 2 to 4 were measured, and the average Arm value minus the background value from a region outside the embryo was defined as the Wg stripe Arm value. In the adjacent interstripe regions we used the same box size to measure and calculate Interstripe Arm values. We also measured the relative difference in Arm accumulation between the Wg Stripes and Interstripes. We also measured the GFP fluorescence of the same boxed regions, allowing us to determine the GFP expression level in different embryos—at times this was used to infer possible genotypes.

### Statistics

Wg-Stripe and Interstripe Arm level values were generally normally distributed, as tested by the D’Agostino-Pearson omnibus normality test as well as the Shapiro-Wilk normality test, and thus parametric tests were employed in statistical analysis. The Paired t-test was used to determine the significance between intragroup values, and an unpaired t-test was used to determine the significance between intergroup values. For multiple comparisons, ordinary one-way ANOVA followed by Dunnett’s multiple comparisons test were applied.

### Immunoblotting

4-8hr old embryos were collected in 0.1% Trition-X100, dechorionated in 50% bleach, and then homogenized with a pestle in RIPA buffer (1% NP-40, 0.5% Na deoxycholate, 0.1% SDS, 50mM Tris pH 8, 300 mM NaCl). Protein concentrations were calculated using Protein Assay Dye (BioRad) following te manufacturer’s recommendations. Samples were mixed with SDS-PAGE sample buffer, boiled for 5 minutes and then run on an 8% SDS-PAGE gel and transferred to a nitrocellulose membrane. Westerns were visualized using a CLX Licor machine which allowed imaged to blots to be collected in a 4-log range. Band densitometry was calculated using LICOR Image Studio and significance was based on a one-sample *t* test using GraphPad. Primary Antibodies: anti-GFP (JL-8 Clontech, mouse monoclonal 1:1000), anti-Axin (a kind gift from Y. Ahmed, guinea pig polyclonal 1:1000), anti-γ-tubulin (Sigma-Aldrich, mouse monoclonal, 1:2000), anti-APC2 (Rabbit polyclonal, a kind gift of M. Bienz [49], 1:1000). Secondary Antibodies: IRDye680RD anti-Rabbit (Licor, 1:10,000), IRDye680RD anti-Guinea pig (Licor, 1:10,000), and IRDye800CW anti-Mouse (Licor 1:10,000).

### RNA-Seq

mRNA collection and RNAseq analysis were as in [50](GEO accession number GSE38727).

### Cell culture and transfections

SW480 cells were cultured at 37° C at normal atmospheric levels of CO_2_ in L15-media (Cellgro) supplemented with 10% FBS and 1x penicillin–streptomycin. *Drosophila* APC2 or Axin protein constructs were transfected into SW480s using Lipofectamine 2000 (Invitrogen) as recommended by the manufacturer. Cells were imaged for yeast fluorescence comparison experiments 24 hours later. Full length *Drosophila* APC2 and Axin were cloned with either a GFP or RFP tag as in [5].

### Yeast Fluorescence Comparison

Yeast Fluorescence comparison analysis was performed as described in [35,36]. Images were taken on a Zeiss LSM 710 confocal microscope, using a 488 diode for stable illumination on a 100x/1.4 NA objective lens. Images were analyzed using FIJI. Yeast strains used for our comparisons to SW480 cells and embryos were Ndc80:GFP (∼306 molecules of GFP) and Mif2:GFP (∼58 molecules of GFP), both were kind gifts from K. Bloom [35]. Briefly, yeast were grown at 24°C in YPD media until they reached an OD between 400-600. Yeast cells were then pelleted and resuspended in YC complete for imaging. Data were used that were consistent with both the Ndc80 and Mif2 standards. Both transfected cells and embryos expressing UAS-Axin:GFP under various GAL4 promoters were imaged live in YC complete media. Each slide was imaged for no longer than 20 minutes at room temperature (∼25°C).

## Author contributions

K.N. Schaefer and M. Peifer conceived the study. D.J. McKay analyzed RNAseq data, T.T. Bonello analyzed protein levels of Axin and APC2, C.E. Williams analyzed Engrailed expression and helped with analysis of lethality and cell fates, and M. Peifer helped with fly husbandry. All other experiments were carried out by K.N. Schaefer and S. Zhang. K.N. Schaefer, S. Zhang, and M. Peifer wrote the manuscript with input from the other authors.

## Acknowledgements

Thanks to M. Bienz, Y. Ahmed, and K. Bloom for key reagents, E. Metzger for lab management and help with fly husbandry, T. Perdue for help with confocal and superresolution microscopy, J. Lawrimore for yeast analysis advice, J. Poulton for statistical advice, and D. Roberts, A. Spracklen and lab members for helpful advice and comments on the manuscript.

## Funding Statement

This work was supported by NIH R35 GM118096 to M.P. K.N.S. was supported by NSF Graduate Fellowship DGE-1650116. The funders had no role in study design, data collection and analysis, decision to publish, or preparation of the manuscript

## Supplemental Figures

**Figure S1:**
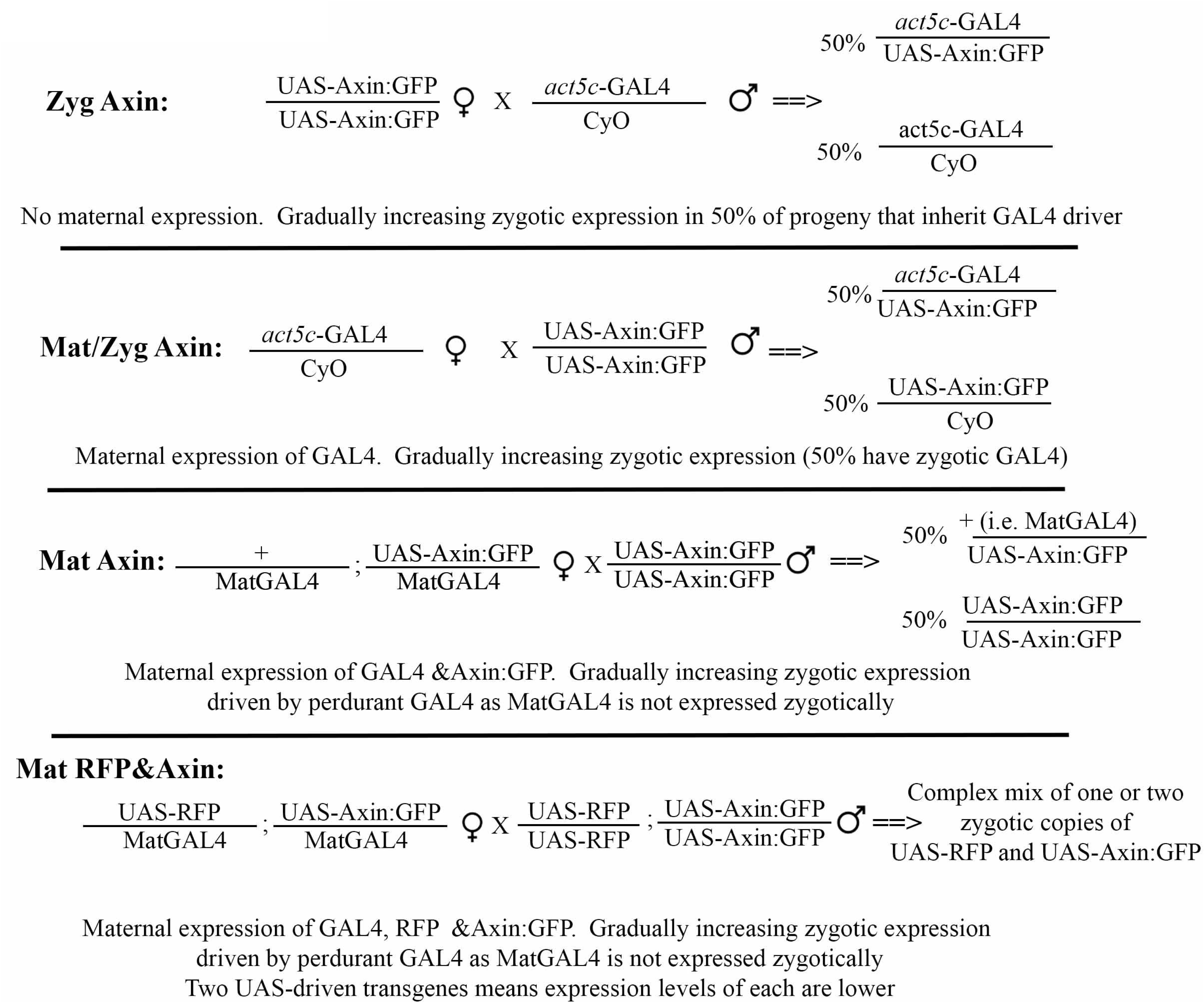
Crosses used to achieve different level and timing of Axin elevation.

**Figure S2.**
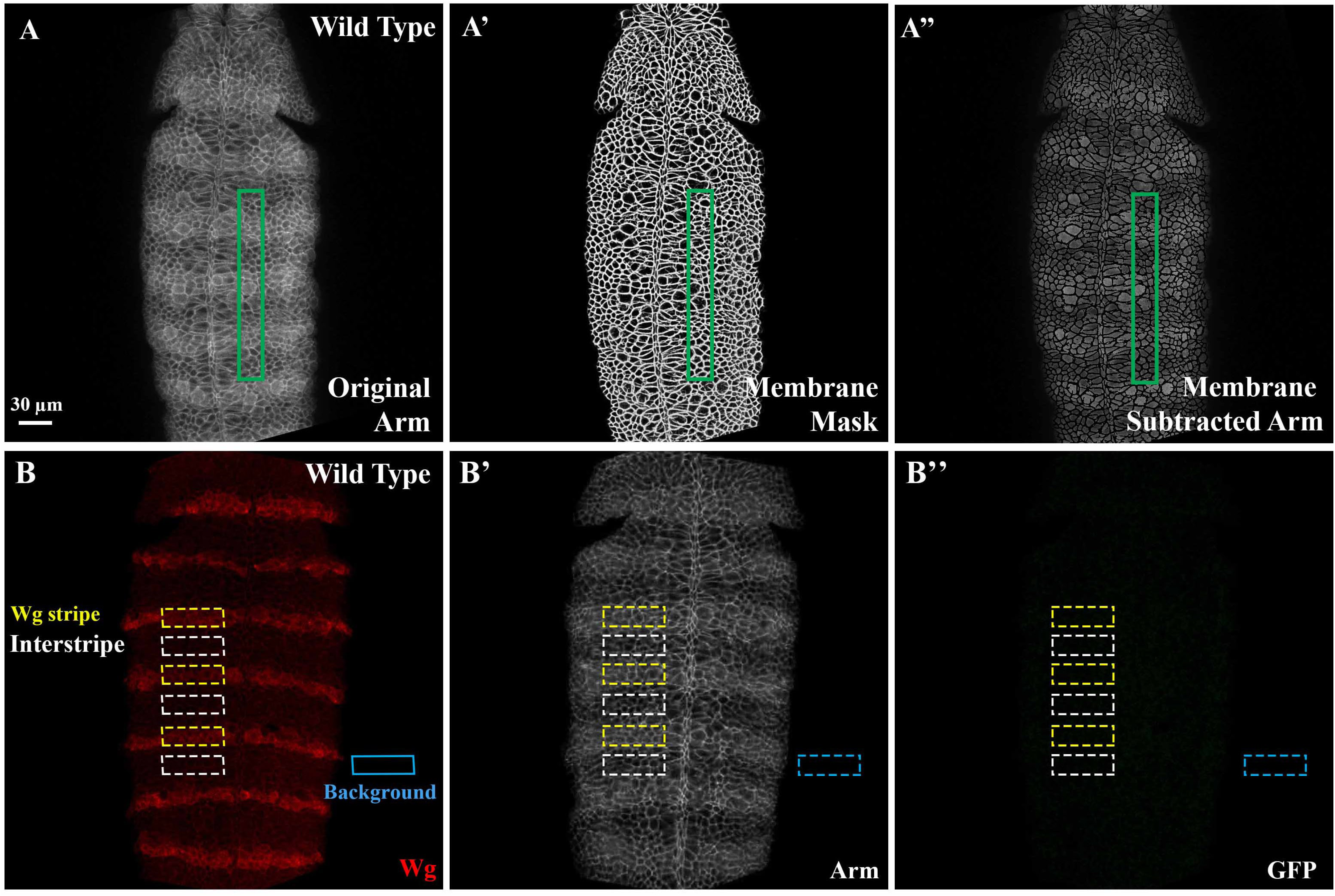
Assessing gradation of Arm levels across the segment and absolute levels of Arm in Wg stripes and interstripes. (A-A”) Representative example illustrating assessment of graded cytoplasmic and nuclear Arm levels across 2-3 embryonic segments (A) Original image of Arm (A’) Plasma membrane mask created using pTyr staining of the same embryo in A. (A”) Resulting Arm image after subtracting A’ from A. The green box illustrates a region of interest (ROI) selected. A profile of the ROI was plotted along the anterior-posterior axis. ROI profiles were adjusted for embryo length and to bring valleys to zero. (B-B”) Representative example illustrating calculations of absolute levels of Arm in Wg expressing cells (Wg-stripes) and Wg-OFF cells (interstripes). Yellow boxes indicate regions sampled for Wg stripes. White boxes represent the regions sampled for the interstripe region. The blue box represents the region sampled for background. In all cases, wildtype and mutant embryos were imaged together using constant imaging conditions.

**Figure S3:**
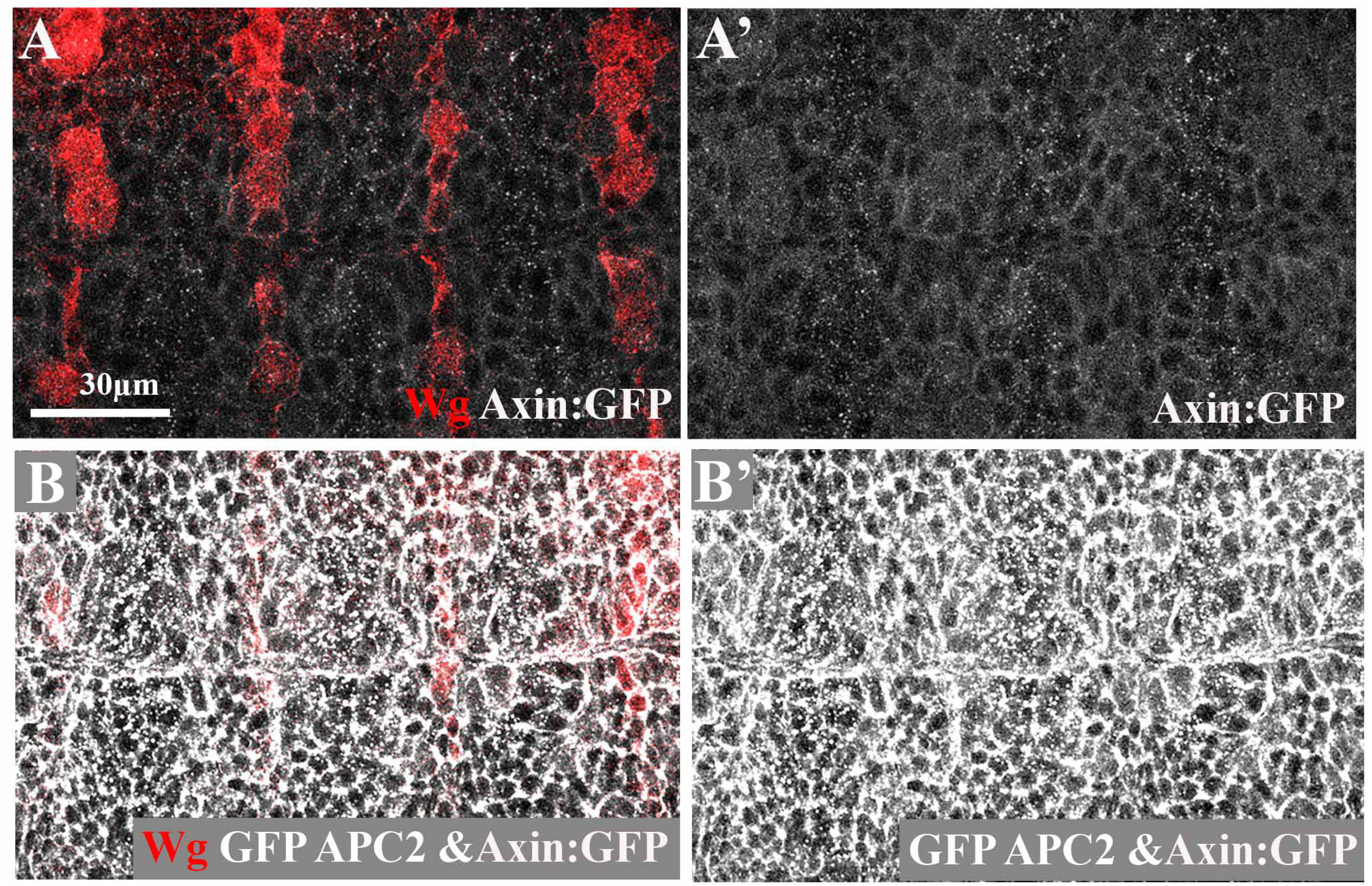
Illustration of how embryos were sorted as to inferred genotype. (A,B) Fixed images of stage 9 embryos imaged under the same microscope conditions, illustrating that GFP:APC2 is substantially brighter under our conditions than Axin:GFP. This is illustrated by the difference in GFP brightness of Mat Axin embryos versus the MatAPC2 & Axin embryos we scored in the GFP high category.

**Figure S4.**
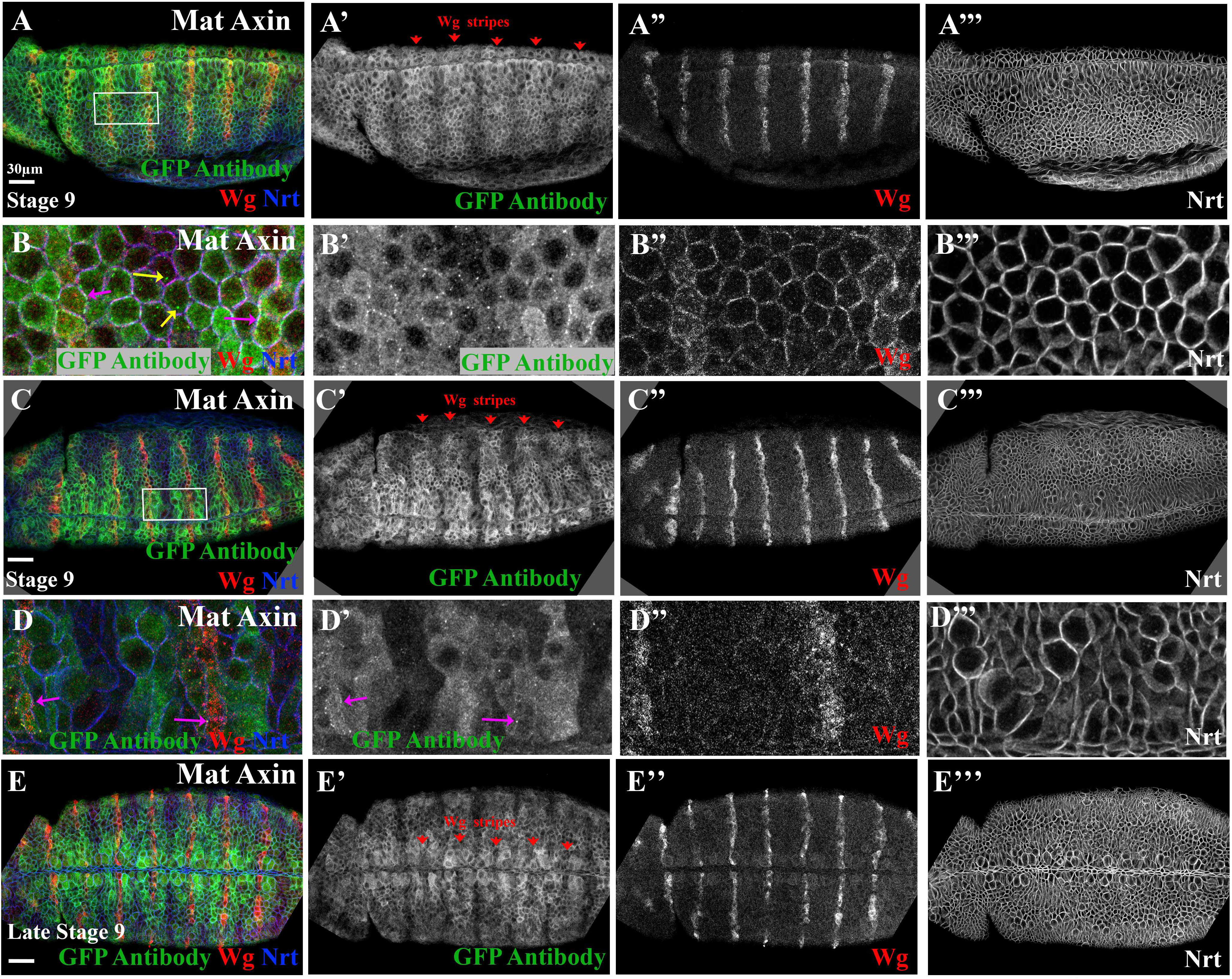
Axin localization using an antibody to the GFP epitope-tag emphasizes changes in cytoplasmic Axin and de-emphasizes Axin puncta in Wg-off cells. (A-D). Stage 9 embryos, anterior to the left. (E) late stage 9-stage 10 embryo. All expressing Axin:GFP using the matGAL4 driver and all stained with antibodies to GFP, Wg, and Neurotactin (Nrt) to visualize plasma membrane. B and D are close-ups of A and C. (A,C) Antibody staining clearly reveals elevated cytoplasmic Axin:GFP in cells receiving Wg signal (arrows). (B) In optimally stained embryos close-ups also reveal both cytoplasmic puncta in Wg-OFF cells and membrane-associated puncta in Wg-ON cells. (D) In many embryos cytoplasmic puncta in Wnt-OFF cells are less apparent, though membrane-associated puncta in Wg-ON cells are visible (arrows). (E). By late stage 9-early stage 10, cells expressing Wg start to accumulate lower levels of cytoplasmic Axin (arrows).

## Supplemental Tables

**Table S1:** Normalized densitometry values. Quantification of protein levels of endogenous and GFP-tagged proteins via immunoblotting followed by band quantification using the CLX Licor, which allows samples to be quantified in a 4-log range. Crosses and their abbreviation are labelled. Means, standard deviation, and number of blots quantified are indicated. Significance was calculated using a one-sample *t-test*.

| Cross (Female x Male) | Abbreviation | Antibody | Mean | Std. Dev. | Significant from Wt<sup>+</sup> | n= |
| --- | --- | --- | --- | --- | --- | --- |
| *act5cGAL4/Cyo* | WT | - | 1.0 | - | - | - |
| *UAS-Axin:GFP x act5cGAL4/Cyo* | Zyg Axin | Axin | 1.0 | 0.5 | No | 10 |
| *UAS-RFP/MatGAL4; UAS-Axin:GFP/MatGAL4 x UAS-RFP; UAS-Axin:GFP* | Mat RFP & Axin | Axin | 3.1 | 1.0 | ** | 5 |
| *actc5cGAL4/Cyo x UAS-Axin:GFP* | Mat/Zyg Axin | Axin | 7.4 | 3.0 | * | 4 |
| *+/MatGAL4; UAS-Axin:GFP/MatGAL4 x UAS-Axin:GFP* | Mat Axin | Axin | 7.9 | 3.0 | * | 4 |
| *APC2>GFP:APC2; APC2<sup>g10</sup>* | GFP:APC2 | APC2 | 0.9 | 0.4 | No | 4 |
| *APC2>GFP:APC2; APC2<sup>g10</sup>* | GFP:APC2 | GFP<sup>#</sup> | 4.3 | 1.4 | *** | 8 |
| *UAS-GFP:APC2/MatGAL4; +/MatGAL4 x UAS-GFP:APC2* | Mat APC2 | GFP<sup>##</sup> | 10.9 | 0.4 | *** | 4 |
| *UAS-GFP:APC2/MatGAL4; UAS-Axin:GFP/MatGAL4 x UAS-GFP:APC2; UAS-Axin:GFP* | Mat APC2 & Axin | Axin | 2.9 | 1.4 | ** | 9 |
| *UAS-GFP:APC2/MatGAL4; UAS-Axin:GFP/MatGAL4 x UAS-GFP:APC2; UAS-Axin:GFP* | Mat APC2 & Axin | GFP<sup>##</sup> | 20.8 | 1.7 | ** | 3 |

+ = significance calculated using one-sample student *t* test; \* = p value < 0.05, \*\* = p value < 0.005, \*\*\* = p value < 0.0005;

**Table S2:** Embryonic viability. Quantification of embryonic viability after altering levels of GFP:APC2 and/or Axin:GFP. Crosses, embryonic viability, standard deviation and numbers of embryos assayed are indicated.

| Cross (Female x Male) | Abbreviation | Viability | StdDev | n= |
| --- | --- | --- | --- | --- |
| *UAS-Axin:GFP x UAS-Axin:GFP* | UAS-Axin:GFP Only | 96.7 | 1.5 | 243 |
| *UAS-Axin:GFP x act5cGAL4/Cyo* | Zyg Axin | 95.4 | 3.5 | 409 |
| *UAS-RFP/MatGAL4; UAS-Axin:GFP/MatGAL4 x UAS-RFP; UAS-Axin:GFP* | Mat RFP & Axin | 68.1 | 6.3 | 568 |
| *+ /MatGAL4; UAS-Axin:GFP/MatGAL4 x UAS-Axin:GFP* | Mat Axin | 22.2 | 2.3 | 900 |
| *actc5cGAL4/Cyo x UAS-Axin:GFP* | Mat/Zyg Axin:GFP | 9.5 | 8.0 | 275 |
| *UAS-GFP:APC2/MatGAL4; + /MatGAL4 x UAS-GFP:APC2* | Mat APC2 | 94.5 | 4.6 | 451 |
| *UAS-GFP:APC2/MatGAL4; UAS-Axin:GFP/MatGAL4 x UAS-GFP:APC2; UAS-Axin:GFP* | Mat APC2 & Axin | 36.9 | 3.9 | 515 |
| *UAS-GFP:APC2/MatGAL4; UAS-Axin:GFP/MatGAL4 x UAS-GFP:APC2* | APC2>>Axin | 79.5 | 7.3 | 667 |
| *UAS-GFP:APC2/MatGAL4; UAS-Axin:GFP/MatGAL4 x UAS-Axin:GFP* | Axin>>APC2 | 29.3 | 6.1 | 791 |

**Table S3:** Embryonic and first instar larva cuticle phenotype. Effects of GFP:APC2 and/or Axin:GFP manipulations on embryonic and first instar larva cuticle phenotypes. Figure 3C shows the cuticle categories. n= number of embryos scored.

| Cross (Female x Male) | Abbreviation | % in Category 1 | % in Category 2 | % in Category 3 | % in Category 4 | % in Category 5 | n= |
| --- | --- | --- | --- | --- | --- | --- | --- |
| *UAS-Axin:GFP x act5cGAL4/Cyo* | Zyg Axin | 75.1 | 18.3 | 6.2 | 0.4 | 0.0 | 149 |
| *UAS-RFP/MatGAL4; UAS-Axin:GFP/MatGAL4 x UAS-RFP; UAS-Axin:GFP* | Mat RFP & Axin | 62.5 | 18.7 | 6.2 | 11.7 | 1.0 | 368 |
| *actc5cGAL4/Cyo x UAS-Axin:GFP* | Mat/Zyg Axin | 10.0 | 10.5 | 13.0 | 19.1 | 47.5 | 323 |
| *+/MatGAL4; UAS-Axin:GFP/MatGAL4 x UAS-Axin:GFP* | Mat Axin | 26.9 | 9.5 | 12.6 | 17.3 | 33.7 | 524 |
| *UAS-GFP:APC2/MatGAL4; +/MatGAL4 x UAS-GFP:APC2* | Mat APC2 | 82.7 | 15.4 | 1.2 | 0.2 | 0.4 | 122 |
| *UAS-GFP:APC2/MatGAL4; UAS-Axin:GFP/MatGAL4 x UAS-GFP:APC2; UAS-Axin:GFP* | Mat APC2 & Axin | 29.0 | 14.8 | 4.7 | 24.9 | 26.7 | 329 |
| *UAS-GFP:APC2/MatGAL4; UAS-Axin:GFP/MatGAL4 x UAS-GFP:APC2* | APC2>>Axin | 51.6 | 35.9 | 5.5 | 2.5 | 4.5 | 128 |
| *UAS-GFP:APC2/MatGAL4; UAS-Axin:GFP/MatGAL4 x UAS-Axin:GFP* | Axin>>APC2 | 16.2 | 21.3 | 12.7 | 10.7 | 39.1 | 635 |

**Table S4:** Rows of En-expressing cells per segment. Quantification of the number of rows of En expressing cells per segment, in embryos in which APC2 or Axin levels are decreased or elevated. n=number of embryos scored. Significance was determined using one-way ANOVA with GraphPad.

| Cross (Female x Male) | Abbreviation | Avg. # of cell rows with En | Std. Dev. | Significant from wt<sup>+</sup> | n= |
| --- | --- | --- | --- | --- | --- |
| *act5cGAL4/Cyo* | WT | 1.8 | 0.2 | - | 21 |
| *UAS-Axin-RNAi/MatGAL4; MatGAL4 x UAS-Axin-RNAi/MatGAL4; MatGAL4* | Axin RNAi | 2.7 | 0.3 | * | 5 |
| *APC2<sup>g10</sup> (APC2 null)* | APC2<sup>g10</sup> | 2.9 | 0.6 | **** | 3 |
| *wg<sup>IG22</sup> /+ (wg null)* | wg<sup>IG22</sup> | 0.5 | 0.5 | **** | 14 |
| *UAS-Axin:GFP x act5cGAL4/Cyo* | Zyg Axin | 1.9 | 0.3 | n.s. | 6 |
| *UAS-RFP/MatGAL4; UAS-Axin:GFP/MatGAL4 x UAS-RFP; UAS-Axin:GFP* | Mat RFP & Axin | 1.5 | 0.3 | n.s. | 12 |
| *+/MatGAL4; UAS-Axin:GFP/MatGAL4 x UAS-Axin:GFP* | Mat Axin | 1.3 | 0.4 | * | 22 |
| *UAS-GFP:APC2/MatGAL4; +/MatGAL4 x UAS-GFP:APC2* | Mat APC2 | 2.2 | 0.2 | n.s. | 10 |

<sup>+</sup> Significance was calculated using one-way ANOVA; n.s. = non-significant; \* = p<.05; \*\*\*\* = p<.0001

**Table S5:** Effects on Arm levels of elevating Axin and/or APC2 levels. Arm accumulation levels in Wg-expressing stripes versus Arm levels in the interstripes for the indicated genotypes as in Figure S2B. These are the raw data used to create the box-and-whisker plots in the Figures. n= number of embryos examined.

| Abbrev. Genotype | Category | Mean | Std. Dev. | Minimum | 25<sup>th</sup> Percentile | Median | 75<sup>th</sup> Percentile | Maximum | n= |
| --- | --- | --- | --- | --- | --- | --- | --- | --- | --- |
| Wildtype<br>(*act5cGAL4/Cyo*) | Wg Stripe | 189502 | 57641 | 115308 | 144436 | 165122 | 241961 | 278825 | 10 |
|  | Interstripe | 121351 | 37219 | 71619 | 96433 | 108979 | 151905 | 189887 | 10 |
| Mat Axin | Wg Stripe | 128196 | 40878 | 79708 | 94402 | 122544 | 146699 | 228161 | 11 |
|  | Interstripe | 110603 | 23538 | 78025 | 90643 | 103552 | 131697 | 143606 | 11 |
| Wildtype<br>(*act5cGAL4/Cyo*) | Wg Stripe | 64909 | 12683 | 46595 | 56127 | 63731 | 72185 | 95562 | 21 |
|  | Interstripe | 48737 | 11239 | 34593 | 38754 | 47431 | 53488 | 80570 | 21 |
| Mat RFP & Axin<br>Lower GFP | Wg Stripe | 59219 | 15569 | 31098 | 46050 | 64029 | 72824 | 80381 | 13 |
|  | Interstripe | 45967 | 12967 | 26661 | 36118 | 46419 | 59746 | 64117 | 13 |
| Mat RFP & Axin<br>Higher GFP | Wg Stripe | 39758 | 7491 | 30852 | 33473 | 38051 | 46225 | 54182 | 10 |
|  | Interstripe | 36796 | 5180 | 27238 | 33647 | 36800 | 42773 | 43351 | 10 |
| Wildtype<br>(*act5cGAL4/Cyo*) | Wg Stripe | 239631 | 57740 | 138431 | 209218 | 229983 | 280509 | 333401 | 11 |
|  | Interstripe | 159690 | 49143 | 91128 | 126440 | 150837 | 180870 | 247271 | 11 |
| Mat APC2 | Wg Stripe | 318561 | 58230 | 253471 | 265841 | 309908 | 372713 | 435928 | 11 |
|  | Interstripe | 187333 | 33560 | 145382 | 159956 | 181812 | 201144 | 265900 | 11 |
| Wildtype<br>(*act5cGAL4/Cyo*) | Wg Stripe | 204637 | 45223 | 79176 | 173770 | 207570 | 234588 | 288438 | 36 |
|  | Interstripe | 135451 | 32366 | 58278 | 112824 | 129429 | 152718 | 220457 | 36 |
| Mat APC2 & Axin<br>APC2+; Axin+ | Wg Stripe | 314265 | 47598 | 235343 | 278999 | 310783 | 346017 | 390044 | 11 |
|  | Interstripe | 163378 | 24203 | 119670 | 140780 | 167185 | 180692 | 198515 | 11 |
| Mat APC2 & Axin<br>APC2+; Axin++ | Wg Stripe | 124596 | 12761 | 105843 | 112659 | 128190 | 133803 | 143265 | 8 |
|  | Interstripe | 98061 | 14441 | 76147 | 87797 | 97210 | 111791 | 118670 | 8 |
| Mat APC2 & Axin<br>APC2++; Axin+ | Wg Stripe | 280121 | 104100 | 159805 | 175731 | 304371 | 366095 | 443029 | 9 |
|  | Interstripe | 154319 | 55917 | 76548 | 93721 | 152046 | 203967 | 226113 | 9 |
| Mat APC2 & Axin<br>APC2++; Axin++ | Wg Stripe | 143706 | 32493 | 93466 | 114848 | 155752 | 164689 | 197860 | 12 |
|  | Interstripe | 121121 | 25701 | 81880 | 94435 | 125967 | 143678 | 154060 | 12 |

**Table S6:** Quantification of the difference in Arm levels in Wg-stripe versus interstripe-cells. Quantification of difference in Arm accumulation between the Wg stripes and interstripes within individual embryos. These are the raw data behind the scatter plots in the Figures. n= number of embryos examined.

| Abbrev. Genotype | Mean | Std. Dev. | Minimum | 25<sup>th</sup><br>Percentile | Median | 75<sup>th</sup><br>Percentile | Maximum | n= |
| --- | --- | --- | --- | --- | --- | --- | --- | --- |
| **Wildtype**<br>*(act5cGAL4/Cyo)* | 68151 | 22105 | 43689 | 48003 | 62445 | 89313 | 103911 | 10 |
| **Mat Axin** | 17593 | 23556 | -780.8 | 1683 | 12259 | 20734 | 84555 | 11 |
| **Wildtype**<br>*(act5cGAL4/Cyo)* | 16172 | 3778 | 8073 | 12384 | 16355 | 19396 | 22201 | 21 |
| **Mat RFP & Axin**<br>Lower GFP | 13252 | 5664 | 4103 | 9550 | 13012 | 18619 | 21253 | 13 |
| **Mat RFP & Axin**<br>Higher GFP | 2962 | 3791 | -2263 | -117.5 | 3331 | 4674 | 10831 | 10 |
| **Wildtype**<br>*(act5cGAL4/Cyo)* | 79941 | 15078 | 47303 | 71183 | 82541 | 94366 | 99639 | 11 |
| **Mat APC2** | 131228 | 34523 | 71658 | 112477 | 121318 | 170029 | 179568 | 11 |
| **Wildtype**<br>*(act5cGAL4/Cyo)* | 69186 | 21282 | 20899 | 54668 | 68992 | 82661 | 111353 | 36 |
| **Mat APC2 & Axin**<br>APC2+; Axin+ | 150886 | 40165 | 63303 | 125452 | 156161 | 187565 | 204232 | 11 |
| **Mat APC2 & Axin**<br>APC2+; Axin++ | 26535 | 7269 | 16321 | 19646 | 26388 | 32595 | 37605 | 8 |
| **Mat APC2 & Axin**<br>APC2++; Axin+ | 125801 | 55865 | 60347 | 81289 | 113078 | 176501 | 216917 | 9 |
| **Mat APC2 & Axin**<br>APC2++; Axin++ | 22585 | 10484 | 9084 | 16869 | 20052 | 28007 | 49430 | 12 |

**Table S7:** Fluorescence comparison values. Detailed results from calculating the number of GFP-tagged Axin or APC2 proteins in Axin puncta in live human SW480 colon cancer cells or in *Drosophila* embryo. These data form the basis of Figure 9. n=number of puncta examined.

| Name | Organism | Mean # GFP<br>molc. | Range | Std. Dev. | n= |
| --- | --- | --- | --- | --- | --- |
| Axin:GFP Only | Human Colon Cancer cells<br>(SW480) | 728 | 163-1327 | 333 | 38 |
| Axin:GFP + RFP:APC2 | Human Colon Cancer cells<br>(SW480) | 801 | 104-2041 | 457 | 39 |
| GFP:APC2 + Axin:RFP | Human Colon Cancer cells<br>(SW480) | 917 | 162-3297 | 608 | 55 |
| Mat RFP & Axin:GFP | *Drosophila* Embryo | 195 | 46-931 | 152 | 70 |
| Mat RFP & Axin:GFP<br>**Wg ON** | *Drosophila* Embryo | 128 | 46-305 | 61 | 35 |
| Mat RFP & Axin:GFP<br>**Wg OFF** | *Drosophila* Embryo | 262 | 56-931 | 185 | 35 |

## References

1. Clevers H, Nusse R. Wnt/beta-catenin signaling and disease. Cell. 2012; 149: 1192–1205.

2. Nusse R, Clevers H. Wnt/beta-Catenin Signaling, Disease, and Emerging Therapeutic Modalities. Cell. 2017; 169: 985–999.

3. Kishida S, Yamamoto H, Hino SI, Ikeda S, Kishida M, et al. DIX domains of Dvl and axin are necessary for protein interactions and their ability to regulate beta-catenin stability. Molecular and Cellular Biology. 1999; 19: 4414–4422.

4. Schwarz-Romond T, Fiedler M, Shibata N, Butler PJ, Kikuchi A, et al. The DIX domain of Dishevelled confers Wnt signaling by dynamic polymerization. Nat Struct Mol Biol. 2007; 14: 484–492.

5. Pronobis MI, Rusan NM, Peifer M. A novel GSK3-regulated APC:Axin interaction regulates Wnt signaling by driving a catalytic cycle of efficient betacatenin destruction. Elife. 2015; 4: e08022.

6. Kunttas-Tatli E, Roberts DM, McCartney BM. Self-association of the APC tumor suppressor is required for the assembly, stability, and activity of the Wnt signaling destruction complex. Mol Biol Cell. 2014; 25: 3424–3436.

7. Hernandez AR, Klein AM, Kirschner MW. Kinetic responses of beta-catenin specify the sites of Wnt control. Science. 2012; 338: 1337–1340.

8. Li VS, Ng SS, Boersema PJ, Low TY, Karthaus WR, et al. Wnt Signaling through Inhibition of beta-Catenin Degradation in an Intact Axin1 Complex. Cell. 2012; 149: 1245–1256.

9. Kim SE, Huang H, Zhao M, Zhang X, Zhang A, et al. Wnt stabilization of beta-catenin reveals principles for morphogen receptor-scaffold assemblies. Science. 2013; 340: 867–870.

10. Lee E, Salic A, Kirschner MW. Physiological regulation of {beta}-catenin stability by Tcf3 and CK1{epsilon}. Journal of Cell Biology. 2001; 154: 983–994.

11. Lee E, Salic A, Kruger R, Heinrich R, Kirschner MW. The roles of APC and Axin derived from experimental and theoretical analysis of the Wnt pathway. PLoS Biol. 2003; 1: E10.

12. Tan CW, Gardiner BS, Hirokawa Y, Layton MJ, Smith DW, et al. Wnt signalling pathway parameters for mammalian cells. PLoS One. 2012; 7: e31882.

13. Cliffe A, Hamada F, Bienz M. A role of Dishevelled in relocating Axin to the plasma membrane during wingless signaling. Curr Biol. 2003; 13: 960–966.

14. Wang Z, Tacchelly-Benites O, Yang E, Thorne CA, Nojima H, et al. Wnt/Wingless Pathway Activation Is Promoted by a Critical Threshold of Axin Maintained by the Tumor Suppressor APC and the ADP-Ribose Polymerase Tankyrase. Genetics. 2016; 203: 269–281.

15. Yang E, Tacchelly-Benites O, Wang Z, Randall MP, Tian A, et al. Wnt pathway activation by ADP-ribosylation. Nat Commun. 2016; 7: 11430.

16. Peterson-Nedry W, Erdeniz N, Kremer S, Yu J, Baig-Lewis S, et al. Unexpectedly robust assembly of the Axin destruction complex regulates Wnt/Wg signaling in Drosophila as revealed by analysis in vivo. Dev Biol. 2008; 320: 226–241.

17. Feng Y, Li X, Ray L, Song H, Qu J, et al. The Drosophila tankyrase regulates Wg signaling depending on the concentration of Daxin. Cell Signal. 2014; 26: 1717–1724.

18. MacDonald BT, He X. A finger on the pulse of Wnt receptor signaling. Cell Res. 2012; 22: 1410–1412.

19. Fiedler M, Mendoza-Topaz C, Rutherford TJ, Mieszczanek J, Bienz M. Dishevelled interacts with the DIX domain polymerization interface of Axin to interfere with its function in down-regulating beta-catenin. Proceedings of the National Academy of Sciences of the United States of America. 2011; 108: 1937–1942.

20. Mao J, Wang J, Liu B, Pan W, Farr GH, 3rd, et al. Low-density lipoprotein receptor-related protein-5 binds to Axin and regulates the canonical Wnt signaling pathway. Mol Cell. 2001; 7: 801–809.

21. Tolwinski NS, Wehrli M, Rives A, Erdeniz N, DiNardo S, et al. Wg/Wnt signal can be transmitted through arrow/LRP5,6 and Axin independently of Zw3/Gsk3beta activity. Dev Cell. 2003; 4: 407– 418.

22. Wang Z, Tacchelly-Benites O, Yang E, Ahmed Y. Dual Roles for Membrane Association of Drosophila Axin in Wnt Signaling. PLoS Genet. 2016; 12: e1006494.

23. McCartney BM, Dierick HA, Kirkpatrick C, Moline MM, Baas A, et al. Drosophila APC2 is a cytoskeletally-associated protein that regulates Wingless signaling in the embryonic epidermis. J Cell Biol. 1999; 146: 1303–1318.

24. Akong K, Grevengoed E, Price M, McCartney B, Hayden M, et al. Drosophila APC2 and APC1 Play Overlapping Roles in Wingless Signaling in the Embryo and Imaginal Discs. Dev Biol. 2002; 250: 91–100.

25. Ahmed Y, Nouri A, Wieschaus E. Drosophila Apc1 and Apc2 regulate Wingless transduction throughout development. Development. 2002; 129: 1751–1762.

26. Ahmed Y, Hayashi S, Levine A, Wieschaus E. Regulation of Armadillo by a *Drosophila* APC Inhibits Neuronal Apoptosis during Retinal Development. Cell. 1998; 93: 1171–1182.

27. Akong K, McCartney B, Peifer M. Drosophila APC2 and APC1 Have Overlapping Roles in the Larval Brain Despite Their Distinct Intracellular Localizations. Dev Biol. 2002; 250: 71–90.

28. Rorth P. Gal4 in the Drosophila female germline. Mech Dev. 1998; 78: 113–118.

29. Brand AH, Perrimon N. Targeted gene expression as a means of altering cell fates and generating dominant phenotypes. Development. 1993; 118: 401–415.

30. Roberts DM, Pronobis MI, Poulton JS, Waldmann JD, Stephenson EM, et al. Deconstructing the beta-catenin destruction complex: mechanistic roles for the tumor suppressor APC in regulating Wnt signaling. Mol Biol Cell. 2011; 22: 1845–1863.

31. Bejsovec A, Martinez-Arias A. Roles of *wingless* in patterning the larval epidermis of *Drosophila*. Development. 1991; 113: 471–485.

32. Peifer M, Sweeton D, Casey M, Wieschaus E. *wingless* signal and Zeste-white 3 kinase trigger opposing changes in the intracellular distribution of Armadillo. Development. 1994; 120: 369–380.

33. Faux MC, Coates JL, Catimel B, Cody S, Clayton AH, et al. Recruitment of adenomatous polyposis coli and beta-catenin to axin-puncta. Oncogene. 2008; 27: 5808–5820.

34. Mendoza-Topaz C, Mieszczanek J, Bienz M. The APC tumour suppressor is essential for Axin complex assembly and function, and opposes Axin’s interaction with Dishevelled. Biology Open. 2011; 1.

35. Lawrimore J, Bloom KS, Salmon ED. Point centromeres contain more than a single centromere-specific Cse4 (CENP-A) nucleosome. J Cell Biol. 2011; 195: 573–582.

36. Verdaasdonk JS, Lawrimore J, Bloom K. Determining absolute protein numbers by quantitative fluorescence microscopy. Methods Cell Biol. 2014; 123: 347–365.

37. Aoki K, Taketo MM. Adenomatous polyposis coli (APC): a multi-functional tumor suppressor gene. J Cell Sci. 2007; 120: 3327–3335.

38. Poulton JS, Mu FW, Roberts DM, Peifer M. APC2 and Axin promote mitotic fidelity by facilitating centrosome separation and cytoskeletal regulation. Development. 2013; 140: 4226–4236.

39. Schwarz-Romond T, Metcalfe C, Bienz M. Dynamic recruitment of axin by Dishevelled protein assemblies. J Cell Sci. 2007; 120: 2402–2412.

40. Pronobis MI, Deuitch N, Posham V, Mimori-Kiyosue Y, Peifer M. Reconstituting regulation of the canonical Wnt pathway by engineering a minimal beta-catenin destruction machine. Mol Biol Cell. 2017; 28: 41–53.

41. Nakamura T, Hamada F, Ishidate T, Anai K, Kawahara K, et al. Axin, an inhibitor of the Wnt signalling pathway, interacts with beta-catenin, GSK-3beta and APC and reduces the beta-catenin level. Genes to Cells. 1998; 3: 395–403.

42. Willert K, Logan CY, Arora A, Fish M, Nusse R. A Drosophila Axin homolog, Daxin, inhibits Wnt signaling. Development. 1999; 126: 4165–4173.

43. Takacs CM, Baird JR, Hughes EG, Kent SS, Benchabane H, et al. Dual positive and negative regulation of wingless signaling by adenomatous polyposis coli. Science. 2008; 319: 333–336.

44. Yamulla RJ, Kane EG, Moody AE, Politi KA, Lock NE, et al. Testing models of the APC tumor suppressor/beta-catenin interaction reshapes our view of the destruction complex in Wnt signaling. Genetics. 2014; 197: 1285–1302.

45. McCartney BM, Price MH, Webb RL, Hayden MA, Holot LM, et al. Testing hypotheses for the functions of APC family proteins using null and truncation alleles in Drosophila. Development. 2006; 133: 2407–2418.

46. Wieschaus E, Nüsslein-Volhard C (1986) Looking at embryos. In: Roberts DB, editor. Drosophila, A Practical Approach. Oxford, England: IRL Press. pp. 199–228.

47. Fox DT, Peifer M. Abelson kinase (Abl) and RhoGEF2 regulate actin organization during cell constriction in Drosophila. Development. 2007; 134: 567–578.

48. Peifer M. The product of the Drosophila segment polarity gene armadillo is part of a multi-protein complex resembling the vertebrate adherens junction. Journal of Cell Science. 1993; 105: 993– 1000.

49. Yu X, Waltzer L, Bienz M. A new Drosophila APC homologue associated with adhesive zones of epithelial cells. Nature Cell Biology. 1999; 1: 144–151.

50. McKay DJ, Lieb JD. A common set of DNA regulatory elements shapes Drosophila appendages. Dev Cell. 2013; 27: 306–318.

